# Mapping protein distribution in the canine photoreceptor sensory cilium and calyceal processes by ultrastructure expansion microscopy

**DOI:** 10.1101/2024.06.27.600953

**Authors:** Kei Takahashi, Raghavi Sudharsan, William A. Beltran

## Abstract

Photoreceptors are highly polarized sensory neurons, possessing a unique ciliary structure known as the photoreceptor sensory cilium (PSC). Vertebrates have two subtypes of photoreceptors: rods, which are responsible for night vision, and cones, which support daylight vision and color perception. Despite identifying functional and morphological differences between these subtypes, ultrastructural analyses of the PSC molecular architecture in rods and cones are still lacking. In this study, we employed ultrastructure expansion microscopy (U-ExM) to characterize the molecular architecture of the PSC in canine retina. We demonstrated that U-ExM is applicable to both non-frozen and cryopreserved retinal tissues with standard paraformaldehyde fixation. Using this validated U-ExM protocol, we revealed the molecular localization of numerous ciliopathy-related proteins in canine photoreceptors. Furthermore, we identified significant architectural differences in the PSC, ciliary rootlet, and calyceal processes between canine rods and cones. These findings pave the way for a better understanding of alterations in the molecular architecture of the PSC in canine models of retinal ciliopathies.

## Introduction

Photoreceptors are highly polarized sensory neurons, located in the outermost layer of the neuroretina that play a crucial role in light detection and signal transduction. In vertebrate retinas, there are two subtypes of photoreceptors: rods, which are highly sensitive to light and essential for night vision, and cones, which provide high visual acuity, color perception, and daylight vision [1]. These cells possess a unique sensory ciliary structure known as the photoreceptor sensory cilium (PSC). Within the PSC, photoreception and phototransduction occur in the outer segment (OS), a highly elaborated membrane structure supported by a nine-fold microtubule axoneme that originates from the basal body (BB) in the inner segment (IS). The region connecting the OS and IS, termed the connecting cilium (CC), is characterized by Y-shaped linkers (Y-links) that anchor each microtubule axoneme to the cell membrane and function as a critical gateway for ciliary trafficking [2, 3] (**Figure 1**). These fundamental features of the tubulin-based PSC architecture are thought to be shared between rods and cones, although detailed structural analyses for cone PSC are still lacking [4].

**Figure 1.**
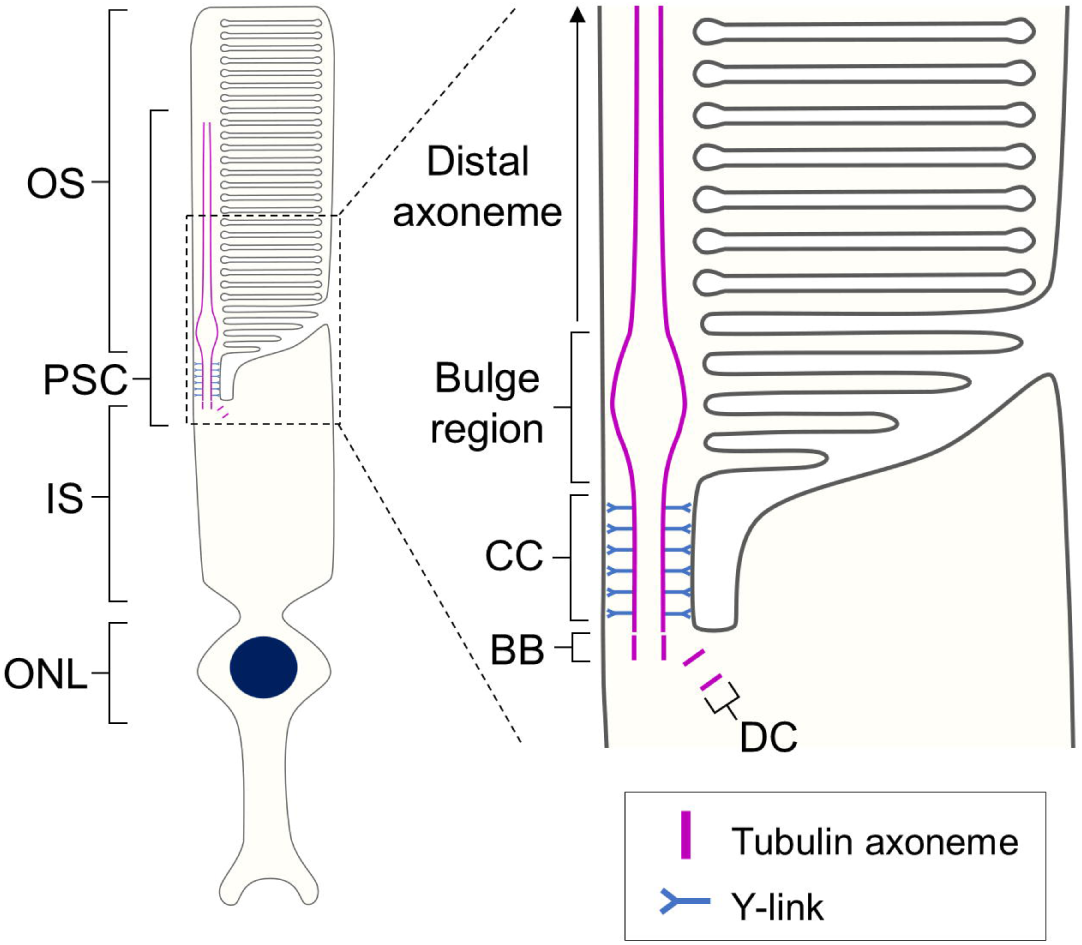
Schematic diagram of the rod photoreceptor sensory cilium. Overall view of the architecture of the rod photoreceptor (left) and detailed depiction of each compartment in the PSC (right). The PSC tubulin axoneme originates from the BB located in the IS and extends through the CC to reach the OS. Just beyond the CC, the axoneme expands at the bulge region and then tapers toward the distal part of the OS. Abbreviations: BB, basal body; CC, connecting cilium; DC, daughter centriole; IS, inner segment; ONL, outer nuclear layer; OS, outer segment; PSC, photoreceptor sensory cilium.

Using cryo-electron tomography, the ultrastructure of rod PSC has been thoroughly analyzed in the mouse retina, revealing that the CC structure of mouse rods is approximately 1.1 μm in length and 300 nm in diameter [5]. Additionally, immunogold electron microscopy has been employed to investigate the precise localization of proteins within this small subcellular compartment, identifying the distribution of several ciliary proteins in mouse rod PSC [6–8]. However, despite the identification of pathogenic variants in over 50 genes associated with non-syndromic retinal ciliopathies [9], the specific roles and precise distribution of these proteins within the PSC have remained largely uncharacterized due to the limited resolution and detection capabilities of conventional fluorescence immunohistochemistry.

With recent advances in super-resolution imaging techniques, it has become feasible to analyze detailed protein localization and morphology within the PSC using fluorescence immunostaining [10–13]. Notably, expansion microscopy (ExM), a sample preparation-based approach in which the specimen is expanded with a swellable hydrogel polymer and allows for ultrastructure observation without the need for specialized microscopy equipment, has become widely accessible to researchers [14]. Numerous modified ExM protocols have been developed to optimize imaging for specific subcellular compartments and sample conditions [15], some of which have been successfully applied to investigate the ultrastructure of photoreceptors [16–18]. Among these advanced methods, ultrastructure expansion microscopy (U-ExM) is particularly notable for its ability to preserve the native intracellular architecture of biological specimens while enabling nanometer-scale observation using standard primary and secondary antibodies used in immunohistochemistry [19]. U-ExM has already made significant contributions to the molecular mapping of the mouse PSC and the detailed assessment of AAV-based gene augmentation therapy in mouse retinal ciliopathy models [18, 20, 21].

Mice are frequently employed in research on PSC morphology and inherited retinal diseases (IRDs), including retinal ciliopathies, due to their ease of handling, low breeding costs, and suitability for gene editing. However, there are significant functional and structural differences between rodent and human photoreceptors, such as very low percentage of cones, the absence of a foveomacular region, and the lack of calyceal processes [22–24]. To address these limitations and enhance the relevance of preclinical animal models, large animals such as dogs, cats, and pigs are proving to be valuable IRDs models. Notably, canine models hold promise for advancing our understanding of the pathogenic mechanisms of IRDs, owing to their fovea-like cone-rich structure [25] and the identification of an increasing number of naturally-occurring genetic mutations linked to human IRDs [26]. Despite this potential, detailed morphological studies on canine photoreceptors remain limited. In this study, we utilize U-ExM to characterize the molecular architecture of the normal adult canine PSC. We validate the U-ExM method for long-term cryopreserved canine retinal samples and provide a comprehensive analysis of the architectural features of the canine PSC. This includes differences in PSC structure between rods and cones, the distribution of retinal ciliopathy-related proteins, and the presence or absence of calyceal processes.

## Results

### U-ExM protocol is effective on non-frozen and cryopreserved canine retinal tissues

To investigate whether U-ExM could be applied to canine retinal tissues, we tested a U-ExM protocol recently adapted for mouse retina [18] on adult canine retinal tissues subjected to different fixation and storage conditions. The short fixation condition (4% paraformaldehyde (PFA) for 15 minutes) is the same as that used for mouse retina in the previous U-ExM study, while a longer fixation protocol (4% PFA for 3 hours followed by 2% PFA for 24 hours) is routinely used by our lab for long-term cryopreservation of canine retinal tissues.

First, we assessed the structural preservation of the photoreceptor OS after expansion under each sample condition. In non-frozen retinas fixed for the short duration, the OS were severely damaged and unevenly swollen (**Figure 2A**). In contrast, the OS structures of both rods and cones were better preserved in the retinas fixed for the longer duration (**Figure 2B**). Similarly, the OS of the cryosection-derived U-ExM samples exhibited relatively better preservation compared to the short-fixed non-frozen retina, although they showed mild damage (**Figure 2C**). Since slight damage to the OS structure was observed even in the unexpanded cryosections (**Figure 2D**), it is likely that the quality of the cryosectioning was reflected in the sample after expansion.

**Figure 2.**
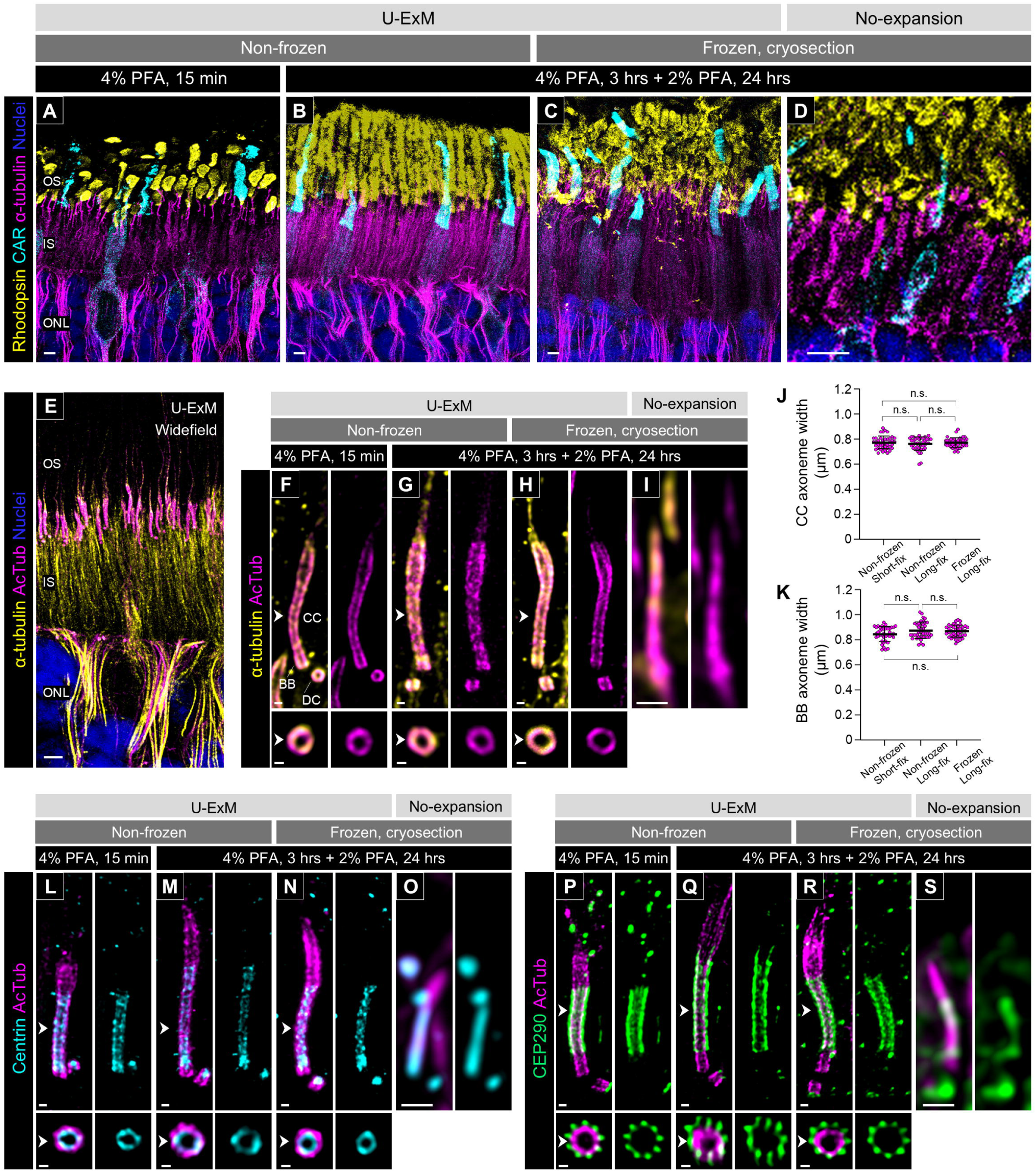
Comparison of tissue integrity and protein localization under different U-ExM conditions. **(A-D)** Low-magnification images of the outer retina stained for rhodopsin (yellow), cone arrestin (CAR; cyan), and α-tubulin (magenta) in U-ExM samples subjected to different preservation/fixation protocols (non-frozen/short-fix, A; non-frozen/long-fix, B; frozen/long-fix, C) and non-expanded samples (D). (D) is enlarged 3.8 times more than (A-C). Scale bars: 5 μm, without correction for expansion factor. **(E)** Widefield view of a U-ExM sample stained for α-tubulin (yellow) and acetylated α-tubulin (AcTub; magenta). Scale bar: 5 μm, without correction for expansion factor. **(F-I)** Details of α-tubulin (yellow) and AcTub (magenta) staining in the canine PSC under U-ExM (F-H) and non-expanded (I) conditions. (I) is enlarged 3.8 times more than (F-H). The lower panels show axial view at the CC for each U-ExM condition, indicated by white arrowheads. Scale bars: lateral view = 500 nm; axial view = 200 nm, without correction for expansion factor. **(J, K)** Widths of AcTub-labeled CC (J) and BB proximal end (K) under different U-ExM conditions. Sample size: n = 40 photoreceptors. For each condition, 4 individual punches or cryosections from 2-4 eyes were stained. Non-significant (n.s., P > 0.05) as assessed by Kruskal-Wallis test with Dunn’s multiple-comparison test. **(L-S)** Molecular localization of Centrin (cyan, L-O) and CEP290 (green, P-S) in normal canine PSC under U-ExM with different preservation/fixation protocols (non-frozen/short-fix, L, P; non-frozen/long-fix, M, Q; frozen/long-fix, N, R) and non-expanded (O, S) conditions. (O, S) are enlarged 3.8 times more than (L-N, P-R). Scale bars: lateral view = 500 nm; axial view = 200 nm, without correction for expansion factor. Abbreviations: BB, basal body; CC, connecting cilium; DC, daughter centriole; INL, inner nuclear layer; IS, inner segment; ONL, outer nuclear layer; OS, outer segment; PFA, paraformaldehyde; PSC, photoreceptor sensory cilium.

Next, to verify whether the expansion factor was affected by fixation duration and storage conditions in canine retinal tissues, we compared the structure of the tubulin axoneme of the PSC under each sample condition. Although non-acetylated tubulin has frequently been used as a PSC axoneme marker in previous U-ExM studies using mouse retina [18, 20, 21], this labeling also showed a distinct signal in the photoreceptor IS. In contrast, immunostaining with antibodies targeting acetylated α-tubulin (AcTub), a widely used marker of primary cilia, labeled the PSC axoneme with high specificity (**Figure 2E**). Furthermore, labeling of acetylated and non-acetylated tubulin at the BB and CC of the PSC showed identical localization in the magnified images of the U-ExM samples, though the non-acetylated tubulin signal was more prominent in the distal axoneme (**Figure 2F-H**). This might suggest that the distal part of the PSC is less prone to acetylation than the BB and CC. We employed AcTub as a highly specific marker for the PSC axoneme in subsequent studies. The width of the AcTub-labeled axoneme on the CC and the proximal end of the BB after processing by U-ExM was comparable across all sample conditions (**Figure 2J and K**). By dividing the CC axoneme width of U-ExM samples by 200.84 nm, which is the previously reported CC axoneme width of mouse photoreceptors [5], we determined an average expansion factor of 3.83 across all sample conditions. Similarly, the gel diameter immediately after the first-round of expansion reached approximately 5 cm in all sample conditions (**Supplementary Figure 2I**), averaging a 4.17 times larger dimension than the original gel diameter. These values are not notably different from the typical expansion factor (∼4) reported in previous U-ExM studies [18, 19].

We further investigated the localization patterns of representative CC-associated proteins to verify whether there are differences in the localization of these proteins among the different sample conditions. Centrin and CEP290 (NPHP6), which have been well characterized in mouse photoreceptors as the CC inner scaffold protein and Y-link-associated protein, respectively [18], were examined. In U-ExM processed canine photoreceptors, Centrin localized specifically inside the tubulin axoneme of the BB and CC (**Figure 2L-N**), while CEP290 was observed outside the CC axoneme (**Figure 2P-R**), consistent with their previously reported localization patterns. Moreover, there were no significant differences in the localization of these proteins among the different sample conditions. In summary, our findings confirmed that U-ExM could be utilized for canine retinal tissue and that variations in PFA fixation and storage conditions do not affect the expansion factor. Considering the better preservation of OS structure, we used long-fixed non-frozen retinas and frozen archival retinal tissues for the subsequent studies described below.

### Differences in tubulin-based molecular architecture between rod and cone PSC

We then analyzed the molecular architecture of the tubulin axoneme in rod and cone PSC of normal adult canine retinas processed by U-ExM. Rod and cone photoreceptors were distinguished by labeling with rhodopsin and cone arrestin (CAR), respectively. The tubulin axoneme in both photoreceptor subtypes was visualized by labeling acetylation and glutamylation on the tubulin, which are well-known post translational modifications in the centriole and primary cilium, using AcTub and GT335 antibodies. In both photoreceptor subtypes, glutamylation was detected on the tubulin axoneme throughout the PSC, similar to acetylation. The glutamylation signal was also found outside the tubulin axoneme only around the CC region (**Figure 3A-D**). A recent study suggested that glutamylation signal shows localization comparable to CEP290 in mouse photoreceptor CC [18]. Similarly, we identified that the region of the glutamylation signal on the tubulin axoneme matches the CEP290 signal on the CC in canine photoreceptors (**Supplementary Figure 3**). Based on these findings, we assumed that the region with glutamylation outside the tubulin axoneme is equivalent to the CC and measured the length and width of each compartment of the PSC axoneme (**Figure 3E**). As previously reported for other cell types [27, 28], the length of the BB was significantly longer than that of the daughter centriole (DC) in each photoreceptor subtype. More interestingly, cone photoreceptors had notably longer DC and BB than rods (Rod DC, 269.71 ± 24.00 μm; Cone DC, 357.27 ± 37.26 μm; Rod BB, 372.06 ± 55.22 μm; Cone BB, 465.84 ± 40.58 μm; **Figure 3F**). In contrast, the length of the CC in cones was significantly shorter than that in rods (Rod CC, 1362.81 ± 164.22 μm; Cone CC, 900.35 ± 169.02 μm; **Figure 3H**). The width of the AcTub-labeled tubulin axoneme in the CC, DC, and BB was identical among the photoreceptor subtypes (**Figure 3G and I**).

**Figure 3.**
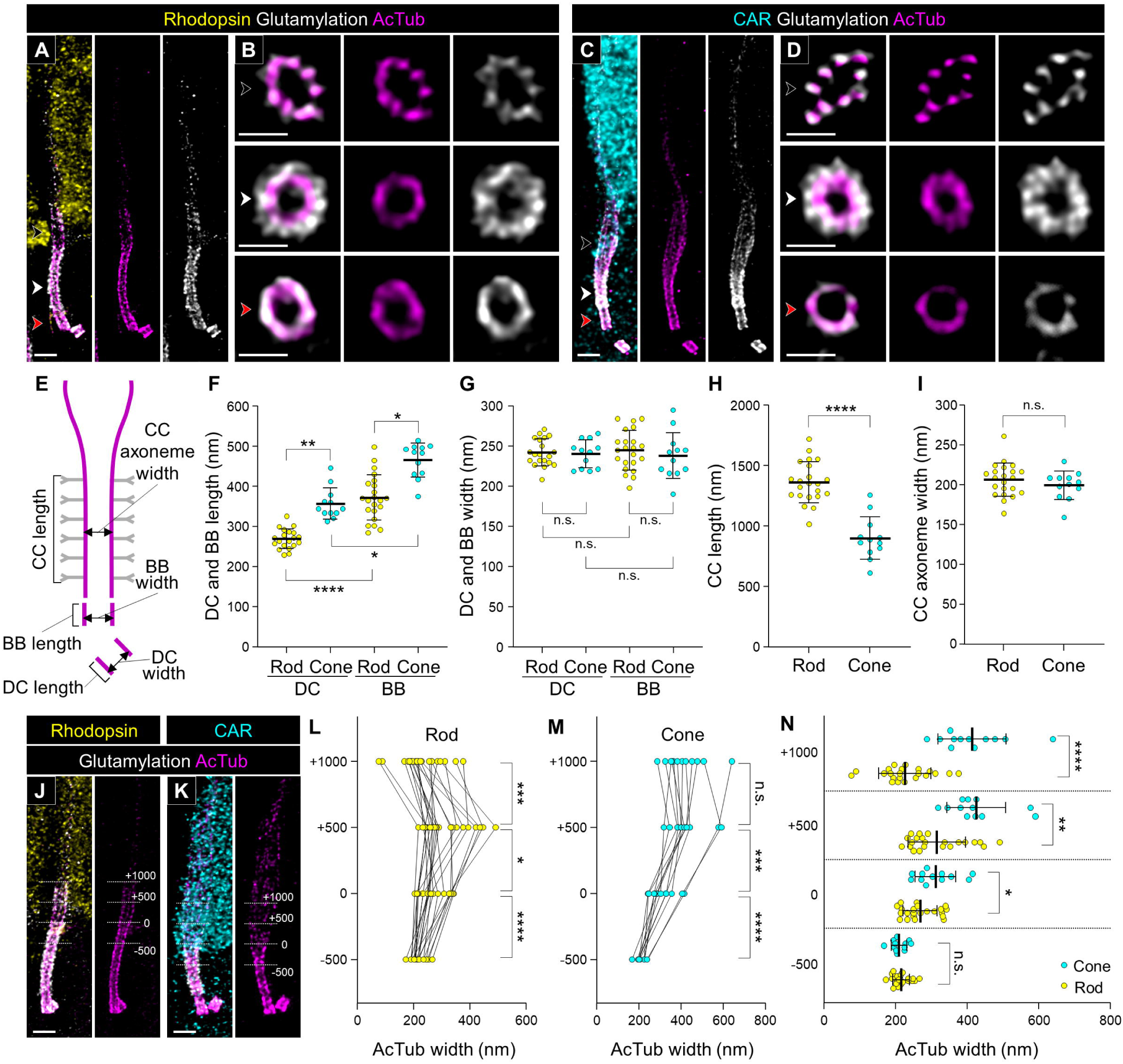
Tubulin-based molecular structure of rod and cone PSC in normal adult canine photoreceptors. **(A-D)** Confocal images of expanded adult canine PSC stained for AcTub (magenta) and glutamylated microtubules (white). Rod and cone photoreceptors were identified by labeling with rhodopsin (yellow, A) and cone arrestin (CAR; cyan, C), respectively. Black, red, and white arrowheads on bulge region, CC, and BB correspond to the respective axial views in (B, D). Scale bars: lateral view = 500 nm; axial view = 200 nm, after correction for expansion factor. **(E)** Schematic diagram showing the measurement of length and width in the CC, BB, and DC. The tubulin axoneme labeled with AcTub and the glutamylation signals on Y-links are represented in magenta and grey, respectively. **(F-I)** Comparison of length and width on the BB (F, G) and the CC (H, I) between rod and cone PSC. **(J, K)** Rod (J) and cone (K) PSC illustrating the measurements of AcTub-labeled axoneme width at four locations relative to the bulge region: +1000, +500, 0, and −500 nm. The distal end of the glutamylation signal located outside of the tubulin axoneme was used to define the 0 nm location. Scale bars: 500 nm, after correction for expansion factor. **(L, M)** Measurements of tubulin width in rod (L) and cone (M) PSC at the four locations illustrated in (J, K). **(N)** Comparison of tubulin width between rod and cone photoreceptors at the four locations shown in (J, K). Sample size: n = 12-22 from ≥ 2 individual eyes. *P < 0.05, **P < 0.01, ***P < 0.001, and ****P < 0.0001 as assessed by Kruskal-Wallis test with Dunn’s multiple-comparison test (F, G) or Mann– Whitney test (H, I, L-N). Abbreviations: BB, basal body; CC, connecting cilium; DC, daughter centriole; PSC, photoreceptor sensory cilium.

Next, we focused on the differences between rod and cone tubulin structures in the outer region of the CC, referred to as the bulge region. Based on the measurement method used in previous studies on mouse photoreceptors [18, 21], we measured the width of the tubulin axoneme at multiple positions along the CC in rod and cone photoreceptors: at the distal end of the CC (0), as well as 500 and 1000 nm above (+500, +1000) and 500 nm below (−500) (**Figure 3J and K**). In the rod PSC, the width of the tubulin axoneme significantly increased from the end of the CC to the +500 nm, then decreased to the +1000 nm (**Figure 3L**). Conversely, in the cone PSC, the enhanced width of tubulin axoneme from the CC endpoint to the +500 nm was maintained even at the +1000 nm (**Figure 3M**). When comparing the axoneme width between rods and cones at each measurement point, the cone axoneme was significantly wider than the rod axoneme at all points except the middle part of the CC (−500) (**Figure 3N**). These results revealed that cone photoreceptors have longer and wider bulge region than rod photoreceptors.

### Molecular mapping of the bulge region and distal axoneme in canine rod and cone PSC

To further characterize the differences in molecular architecture at the distal part of the tubulin axoneme between rods and cones in detail, we investigated the location of several proteins associated with the bulge region and distal axoneme in both rods and cones. LCA5 (lebercilin) has been identified as a bulge region-specific protein in mouse photoreceptors [18, 21]. Consistent with the results from previous studies, LCA5 showed specific localization to the inner side of the bulge region in both rod and cone photoreceptors in the canine PSC (**Figure 4A and B**). Additionally, RP1, known to localize from the bulge region to the distal axoneme in mouse photoreceptors [21], exhibited a comparable localization pattern in canine rod and cone photoreceptors (**Figure 4C and D**). In this study, we identified CCDC66 as a novel protein that specifically localizes to the distal axoneme of both rods and cones (**Figure 4E and F**). CCDC66 is a microtubule-binding protein, and dysfunction of its gene has been reported to cause retinal degeneration in both canine and mouse models [29–31]. While CCDC66 has been shown to localize to the BB and tubulin axoneme in cell lines [32], its distribution patterns in vertebrate retinal tissues have been inconsistent, with reports indicating localization in the OS and IS (**Supplementary Table 5**). In the present study, we tested three commercially available anti-CCDC66 antibodies, including those used in previous studies, on non-expanded canine retinas and found that their reactivity varied significantly (**Supplementary Figure 4A, C, and E**). However, in U-ExM samples, two of the three antibodies (PA5-60642 and sc-102418) produced specific signals on the PSC axoneme (**Supplementary Figure 4B, D, and F**). Although the sc-102418 antibody exhibited more extensive labeling than PA5-60642, both antibodies consistently showed a weak signal near the BB and a prominent signal at the distal axoneme.

**Figure 4.**
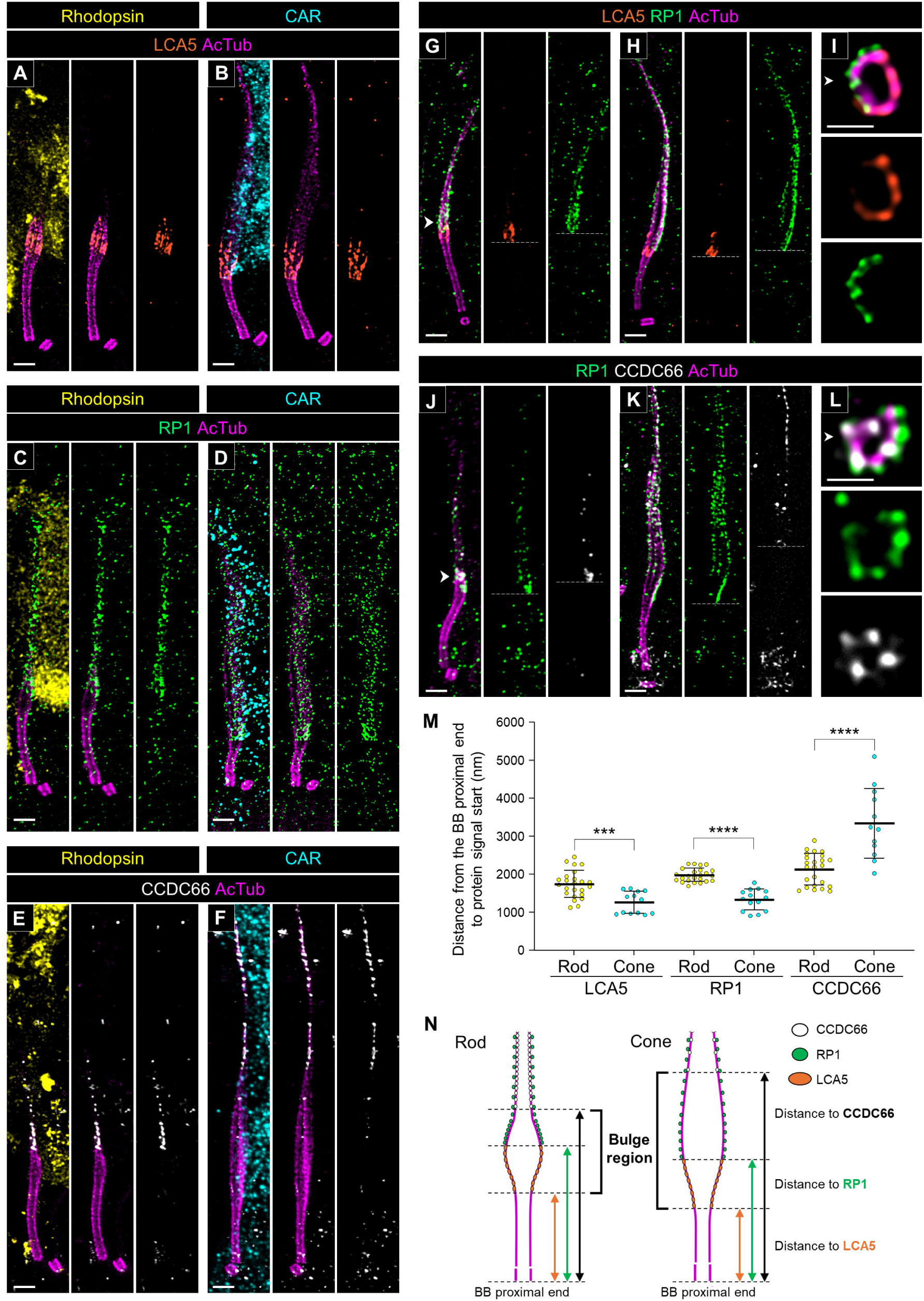
Molecular architecture of the bulge region and distal axoneme in normal canine rod and cone PSC. **(A-F)** Lateral views of expanded adult canine PSC stained for LCA5 (orange, A, B), RP1 (green, C, D), and CCDC66 (white, E, F) in rod (A, C, E) and cone (B, D, F) photoreceptors, respectively. CCDC66 is labeled using an antibody provided by Invitrogen (Cat. #PA5-60642). Scale bars: 500 nm, after correction for expansion factor. **(G-L)** Co-immunolabeling with LCA5/RP1 (orange/green, G-I) and RP1/CCDC66 (green/white, J-L) in each photoreceptor subtype. The right panels (I, L) show axial views of the bulge region indicated by the white arrowheads in each staining. The white dashed lines represent the starting position of each protein signal on the tubulin axoneme. Scale bars: lateral view = 500 nm; axial view = 200 nm, after correction for expansion factor. **(M)** Comparison of distance from the proximal end of the BB to the starting point of each protein signal between rod and cone PSC. Sample size: n = 12-24 from ≥ 3 individual eyes. ***P < 0.001, and ****P < 0.0001 as assessed by Mann–Whitney test. **(N)** Model illustrating the differences in molecular architecture of the distal part of the PSC, including the bulge region, between rod and cone photoreceptors. Abbreviations: BB, basal body; PSC, photoreceptor sensory cilium.

To elucidate the positional relationship of these three proteins in the distal part of the PSC axoneme, we performed co-immunostaining of RP1 with LCA5 and CCDC66, respectively. In both rods and cones, LCA5 localized slightly more basally than RP1, and these two proteins did not co-localize (**Figure 4G and H**). Additionally, axial view images showed that LCA5 localized on the tubulin axoneme while RP1 localized outside the axoneme (**Figure 4I**). CCDC66 appeared to overlap with RP1 in the distal axoneme in the lateral view (**Figure 4J and K**). However, in the axial view, CCDC66 was localized on the tubulin axoneme, distinct from the localized area of RP1 (**Figure 4L**). By measuring the distance from the proximal end of the BB to the starting point of each signal, we found that in cones, LCA5 and RP1 localized more basally compared to rods, while CCDC66 localized more distally (**Figure 4M**). Taken together, LCA5, RP1, and CCDC66 each exhibited unique localization patterns in the distal part of the PSC, and differences in localization of these proteins between rods and cones coincided with the architectural differences in the bulge region between these photoreceptor subtypes (**Figure 4N**).

### Molecular mapping of the connecting cilium in canine rod and cone PSC

Next, we compared the localization of CC-related proteins between rods and cones. POC5, Centrin, and FAM161A have been well-characterized to localize inside the tubulin axoneme as inner scaffold proteins at the BB and CC in mouse photoreceptors [18]. Consistent with the reports in mouse photoreceptors, these three inner scaffold proteins exhibited internal localization within the axoneme at the BB and CC in both canine photoreceptor subtypes (**Figure 5A-I**). Since the endogenous FAM161A signal was too weak to quantify (**Figure 5G-I**), similar to results in mouse photoreceptors [18, 20], we measured the signal length in the CC for only POC5 and Centrin (**Figure 5J**). The signal lengths of these inner scaffold proteins at the CC were significantly shorter in cones than in rods, and these were consistent with the CC lengths measured in Figure 3 (Rod POC5, 1383.09 ± 96.53 μm; Cone POC5, 911.01 ± 129.72 μm; Rod Centrin, 1449.83 ± 115.57 μm; Cone Centrin, 989.27 ± 172.09 μm; **Figure 5K**).

**Figure 5.**
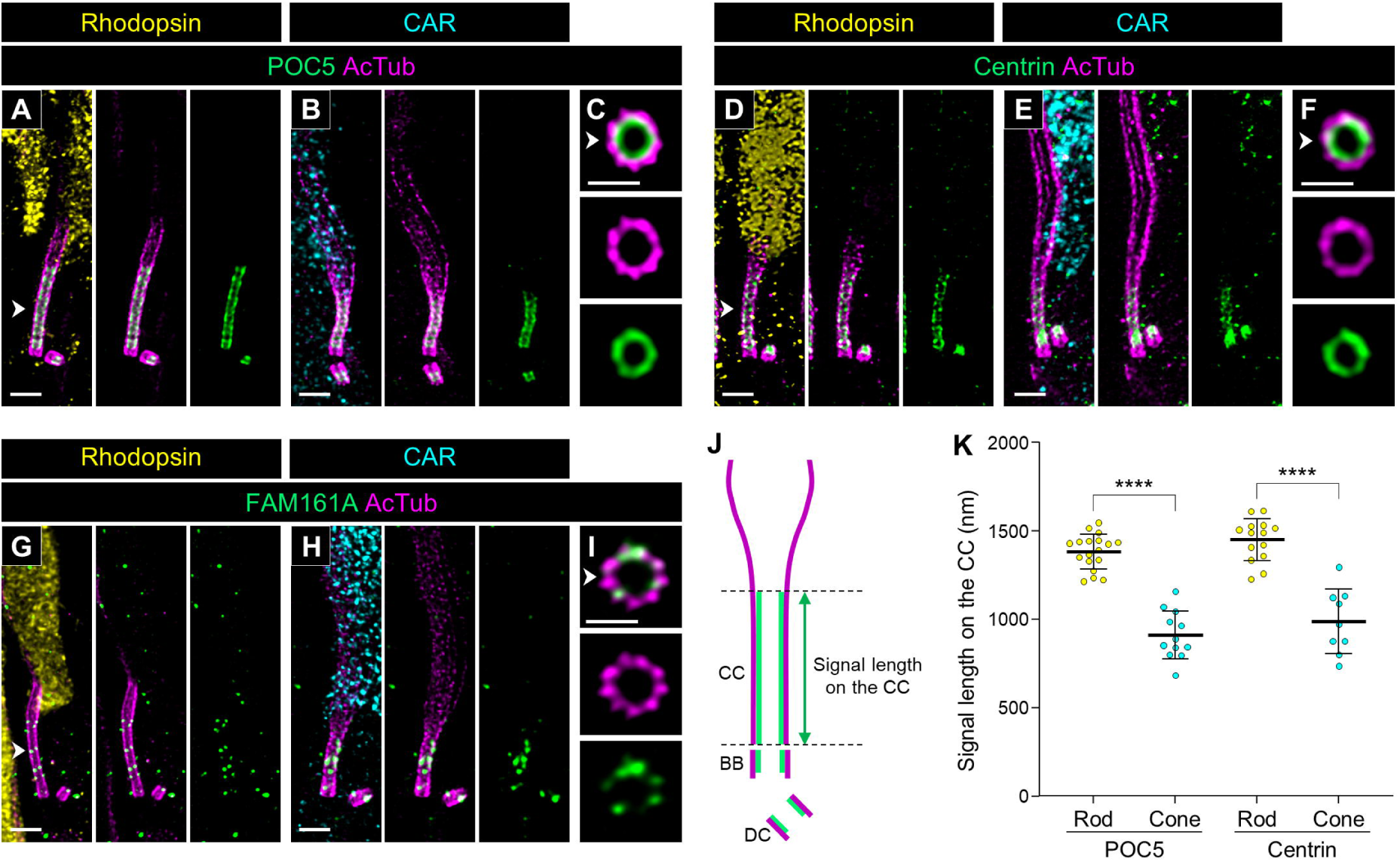
Location of inner scaffold proteins in normal canine rod and cone PSC. **(A-I)** Confocal images of expanded adult canine PSC stained for POC5 (green, A-C), Centrin (green, D-F), and FAM161A (green, G-H). Rod and cone photoreceptors were identified by labeling with rhodopsin (yellow, A, D, G) and CAR (cyan, B, E, H), respectively. The right panels (C, F, I) show axial views of the CC for each protein, indicated by white arrowheads. Scale bars: lateral view = 500 nm; axial view = 200 nm, after correction for expansion factor. **(J)** Schematic diagram illustrating the measurement of signal length in the CC. The tubulin axoneme labeled with AcTub and inner scaffold proteins are represented in magenta and green, respectively. **(K)** Comparison of POC5 and Centrin signal lengths between rod and cone PSC. Sample size: n = 9-18 from ≥ 2 individual eyes. ****P < 0.0001 as assessed by Mann–Whitney test. Abbreviations: CC, connecting cilium; PSC, photoreceptor sensory cilium.

We then investigated the localization of several Y-link-associated proteins in canine photoreceptors. The Y-link is a hallmark element of the transition zone in the primary cilium, exhibiting a distinctive Y-shaped architecture that connects each microtubule doublet to the ciliary membrane [3]. In mammalian photoreceptors, the CC is known as a region corresponding to the transition zone in the primary cilium, and CEP290, a typical Y-link component protein, has been confirmed to localize outside each of the nine tubulin axonemes in the CC of mouse photoreceptors [18]. As shown in Figure 2 and Supplementary Figure 3, CEP290 localized at the CC in canine photoreceptors similar to its localization in mouse photoreceptors. Here, we also found that the signal length of CEP290 at the CC is significantly shorter in cones than in rods, consistent with the lengths of the CC shown in Figure 3 and Figure 5 (Rod CEP290, 1314.33 ± 118.71 μm; Cone CEP290, 891.09 ± 126.21 μm; **Figure 6A, B, C, and S**). Additionally, several other Y-link-associated proteins, including RPGR, RPGRIP1 (LCA6), SPATA7 (LCA3), and NPHP5 (IQCB1), were found to have signal lengths comparable to CEP290 at the CC in both rods and cones (**Figure 6D, E, G, H, J, K, M, N, and S**) and to localize outside the axoneme (**Figure 6F, I, L, O, and R**).

**Figure 6.**
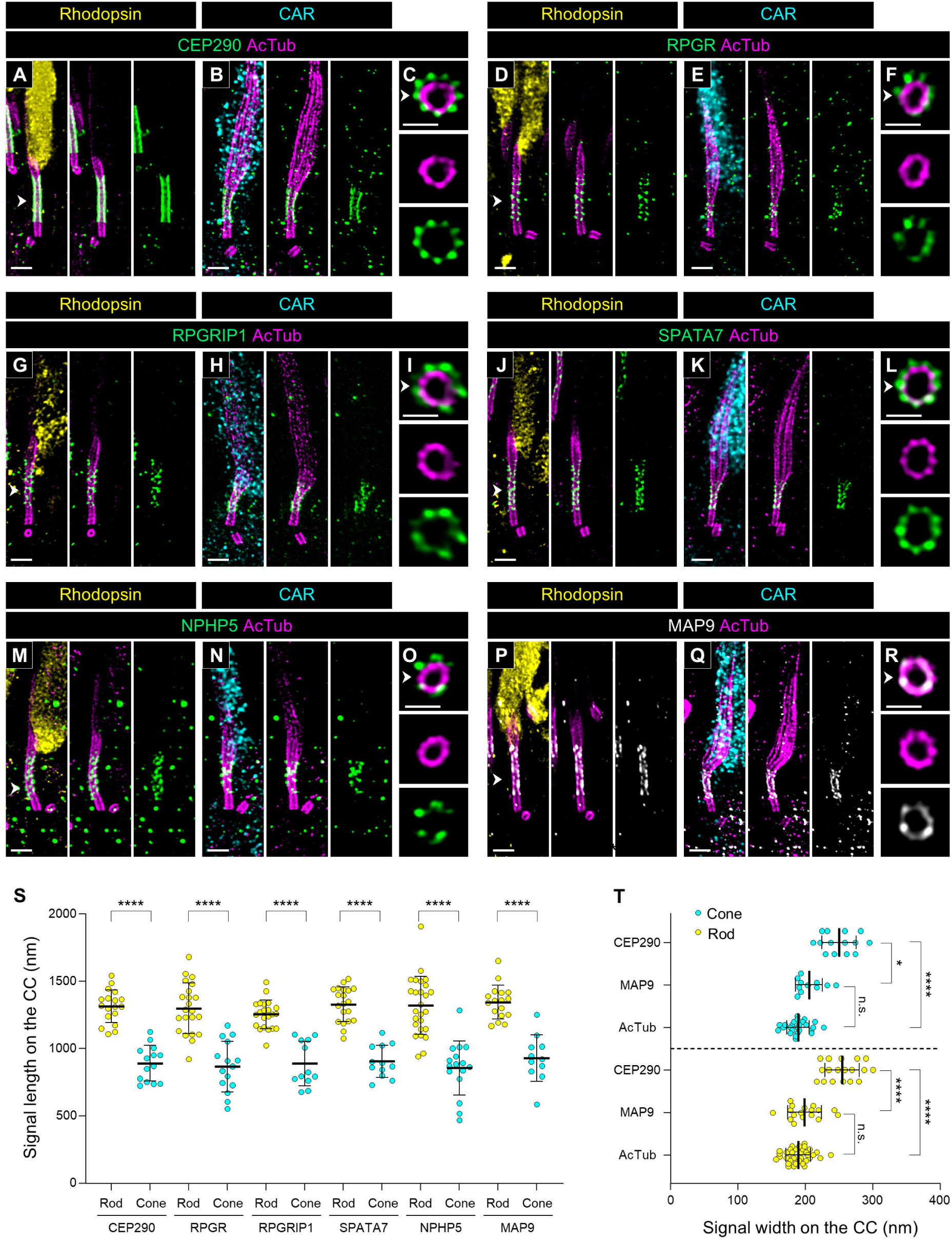
Molecular mapping of multiple connecting cilium-related proteins in normal canine rod and cone PSC. **(A-R)** Expanded normal canine PSC stained for CEP290 (green, A-C), RPGR (green, D-F), RPGRIP1 (green, G-I), SPATA7 (green, J-L), NPHP5 (green, M-O), and MAP9 (white, P-R) in rod (A, D, G, J, M, P) and cone (B, E, H, K, N, Q) photoreceptors, respectively. Axial views of the CC for each protein, indicated by white arrowheads, are shown in (C, F, I, L, O, R). Scale bars: lateral view = 500 nm; axial view = 200 nm, after correction for expansion factor. **(S)** Measurement of signal length of each protein on the CC in rod and cone PSC. Sample size: n = 11-26 from ≥ 3 individual eyes. **(T)** Comparison of signal width between CEP290, MAP9, and AcTub in rod and cone PSC. Sample size: n = 11-35 from ≥ 3 individual eyes. *P < 0.05 and ****P < 0.0001 as assessed by Mann–Whitney test (S) or Kruskal-Wallis test with Dunn’s multiple-comparison test (T). Abbreviations: CC, connecting cilium; PSC, photoreceptor sensory cilium.

Finally, we found that MAP9, a microtubule-binding protein, localizes specifically to the CC in photoreceptors (**Figure 6P and Q**). A homozygous variant in the MAP9 gene has been reported to accelerate disease progression in a naturally-occurring canine *RPGRIP1*-associated cone-rod dystrophy [33, 34]. The co-localization of MAP9 and RPGRIP1 in the CC of both photoreceptor subtypes suggests a direct interaction between these proteins. Additionally, axial view images and quantification of signal width at the CC revealed that MAP9 is located close to the tubulin axoneme, unlike the axial distribution of other Y-link-associated proteins (**Figure 6R and T**). Mapping of these CC-specific proteins confirmed a significant difference in the length of the CC between canine rods and cones.

### Molecular mapping of ciliary trafficking and distal appendage-related proteins on the basal bodies in rod and cone PSC

We next mapped the localization of proteins associated with the BB, the compartment on the basal part of the tubulin-based PSC architecture. The intraflagellar transport (IFT) machinery is responsible for anterograde and retrograde trafficking of ciliary proteins and consists of two multisubunit complexes, IFT-A and IFT-B [35]. In mouse photoreceptors, it was demonstrated using immunoelectron microscopy that several IFT particles have characteristic distribution around the BB and bulge region [8], and similar localization of these particles has recently been confirmed using U-ExM as well [20, 21]. We confirmed that in both canine rods and cones, IFT57, a component of the IFT-B complex [36], has a similar distribution at the BB and bulge region as observed in mouse photoreceptors (**Figure 7A and B**). Additionally, KIF3A, a component of kinesin-II, exhibited a similar distribution to the IFT particles in both photoreceptor subtypes, albeit with a high level of background signal (**Supplementary Figure 5**). Axial view images showed that IFT57 distributes along each of the nine tubulin axonemes at both the BB and bulge region and is located closer to the axonemes at the bulge region than at the BB (**Figure 7C**). We also found that the IFT57 signals at the BB were clearly divided into two layers (**Figure 7A and B**). The distance from the proximal end of the BB to these signals was shorter in rods than in cones (**Figure 7D**), consistent with the difference in BB length between the rods and cones presented in Figure 3.

**Figure 7.**
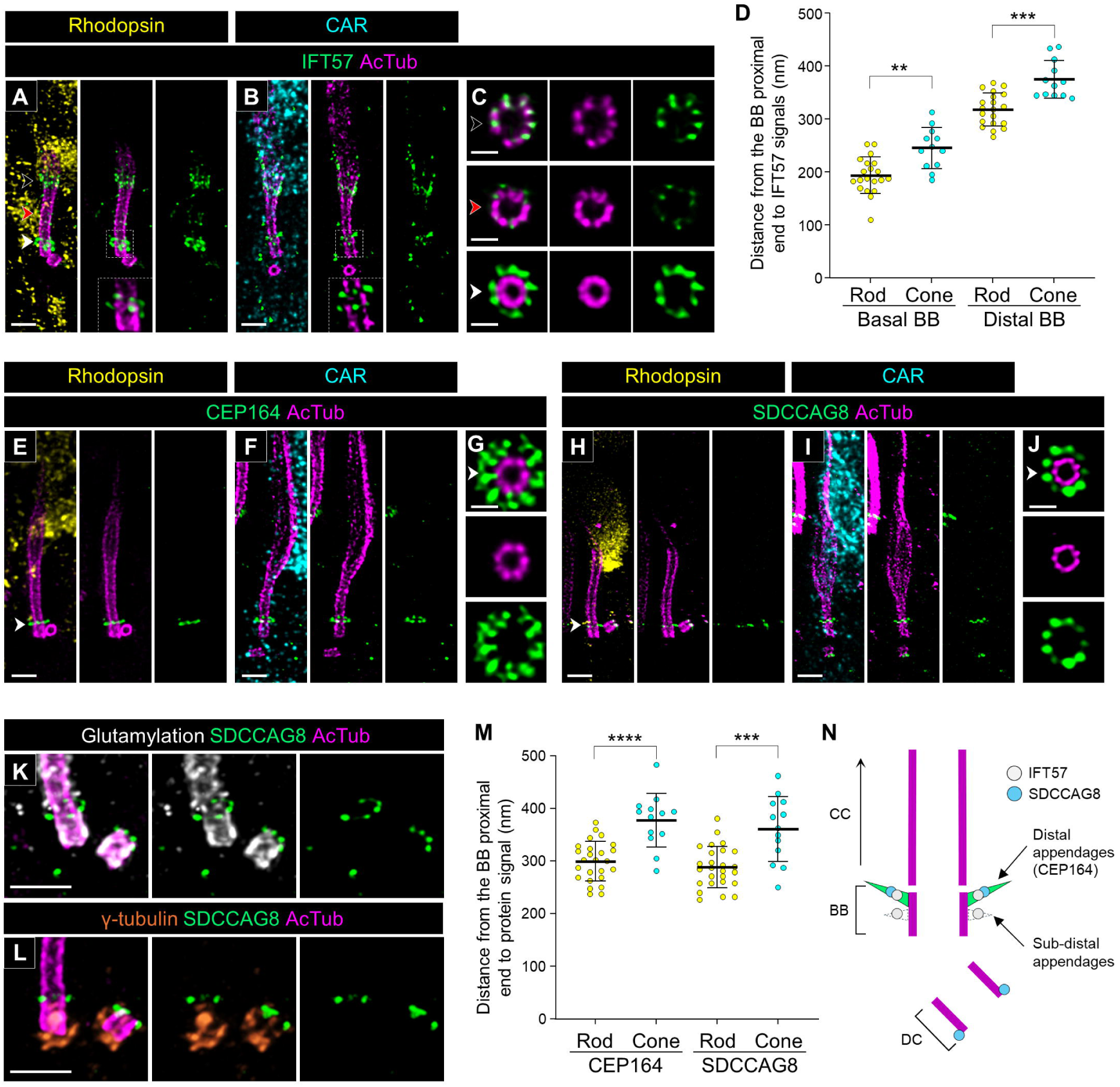
Localization of ciliary trafficking and distal appendage-related proteins in normal canine rod and cone PSC. **(A-C)** Confocal U-ExM images of normal canine PSC stained for AcTub (magenta) and IFT57 (green) in rod (A) and cone (B) photoreceptors, respectively. Insets, which are enlarged single z-scan images from (A) and (B), highlight the detailed localization of IFT57 around the BB. Black, red, and white arrowheads on bulge region, CC, and BB correspond to the axial views shown in (C). Scale bars: lateral view = 500 nm; axial view = 200 nm, after correction for expansion factor. **(D)** Comparison of distance from the proximal end of the BB to IFT57 signals around the basal part of PSC between rod and cone PSC. Sample size: n = 12-20 from ≥ 3 individual eyes. **(E-J)** Expanded normal canine photoreceptors labeled for distal appendage-marker (CEP164; green, E-G) and SDCCAG8 (green, H-J) in rod (E, H) and cone (F, I) photoreceptors, respectively. Axial views of the basal part of PSC for each protein, indicated by white arrowheads, are shown in (G, J). Scale bars: lateral view = 500 nm; axial view = 200 nm, after correction for expansion factor. **(K-L)** Co-immunolabeling with glutamylation/SDCCAG8 (white/green, K) and γ-tubulin/SDCCAG8 (orange/green, L). Scale bars: 500 nm, after correction for expansion factor. **(M)**Measurement of distance from the proximal end of the BB to the signal of each protein in rod and cone PSC. Sample size: n = 13-26 from ≥ 3 individual eyes. **P < 0.01, ***P < 0.001, and ****P < 0.0001 as assessed by Mann–Whitney test. **(N)** Model illustrating the identified location of IFT57 and SDCCAG8 around the distal appendages in normal canine photoreceptors. Abbreviations: BB, basal body; CC, connecting cilium; DC, daughter centriole; PSC, photoreceptor sensory cilium.

Based on the characteristic distribution of IFT57 in the BB, we hypothesized that the IFT molecules localize around distal appendages, also known as transition fibers, in the BB, as has been suggested in previous publications [37, 38]. To verify this, we observed the distribution of CEP164 (NPHP15), a representative distal appendage protein [39], in both rods and cones. As expected, CEP164 was localized in the distal part of the BB in both rods and cones (**Figure 7E and F**), and the axial view exhibited a broader radial distribution (**Figure 7G**), similar to that observed in recent studies using super-resolution microscopy techniques [40, 41]. The distance from the BB proximal end to CEP164 signals was shorter in rods than in cones, and these distances were comparable to these of the IFT57 signals at the distal BB (**Figure 7D and M**).

Additionally, we found that SDCCAG8 (NPHP10), a retinal ciliopathy-associated protein [42, 43], exhibits specific localization to the end of both BB and DC in rods and cones (**Figure 7H and I**). The axial view images at the BB showed that SDCCAG8 has a nine-fold symmetry distribution outside the axoneme, similar to IFT57 and CEP164 (**Figure 7J**). Moreover, co-immunostaining of SDCCAG8 with glutamylation and γ-tubulin demonstrated that the SDCCAG8 signal on the ciliary axoneme is located between the BB and CC (**Figure 7K and L**). The distance from the BB proximal end to SDCCAG8 was comparable to that of CEP164 in both rods and cones (**Figure 7M**). Taken together, these results suggest that IFT57 and SDCCAG8 localize to distal appendages on the BB (**Figure 7N**). Furthermore, based on the distance between distal and subdistal appendages suggested in previous study (∼100 nm), the location of the basal IFT57 signal in the BB is assumed to correspond to the location of the subdistal appendages [44].

### Molecular architecture of the ciliary rootlet in canine photoreceptors

To address the peripheral structures of the PSC in canine photoreceptors, we mapped several proteins related to the ciliary rootlet. Rootletin (CROCC) is a 220-kDa coiled coil protein identified as an architectural component of the rootlet in mouse photoreceptors [45] and has been used as a marker of ciliary rootlet in several studies [46, 47]. Consistent with previous reports, in canine retinas processed with U-ExM, rootletin signals were observed as straight fibers in the photoreceptor IS, representing the architecture of the ciliary rootlets (**Figure 8A**). In the magnified images, a rootlet stem extending straight to the BB and a characteristic branch-like structure extending to the middle of the CC were observed in the rod PSC (**Figure 8B and C, Supplementary Movie 1**). In contrast, cone photoreceptors exhibited a curvilinear rootlet architecture that was quite different from that of rods. In cone photoreceptors, the base of the rootlet did not contact the external limiting membrane, unlike in rods, and the ends of the rootlet appeared to be directed toward the BB and DC, respectively (**Figure 8D and E, Supplementary Movie 2**). These structural features in cones were consistent among cone photoreceptor subtypes, S-cones and L/M-cones (**Supplementary Figure 6**).

**Figure 8.**
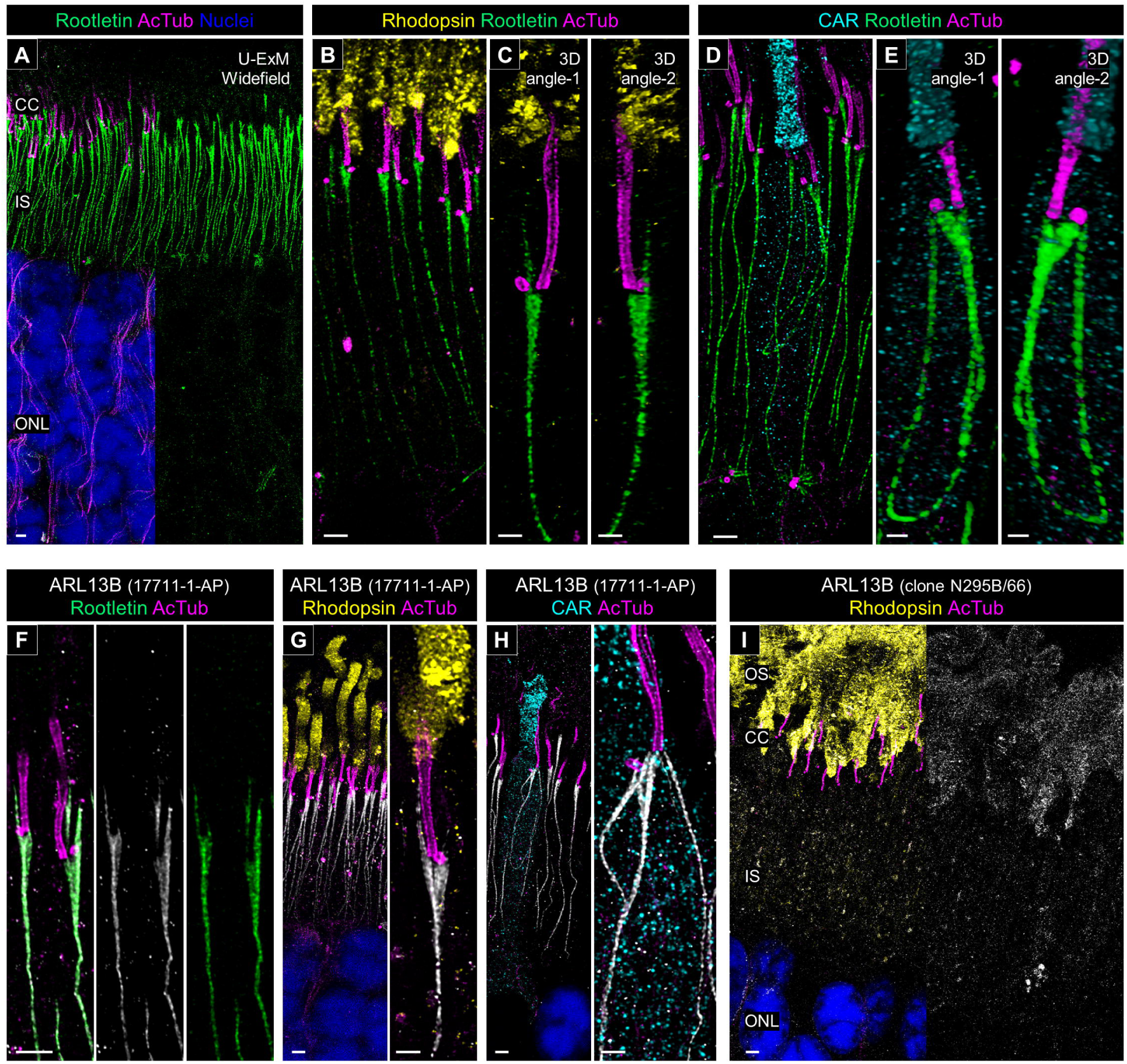
Structural characterization of the ciliary rootlet in normal canine rod and cone photoreceptors. **(A)** Widefield view of a U-ExM sample stained for rootletin (green) and AcTub (magenta). Scale bar: 1 μm, after correction for expansion factor. **(B-E)** High-magnification of expanded normal canine PSC stained for rootletin (green) and AcTub (magenta) in rod (B, C) and cone (D, E) photoreceptors, respectively. 3D rendering images were shown in (C, E) from different angles. Scale bars: 2D maximum projection = 1 μm; 3D view = 500 nm, after correction for expansion factor. **(F)** Co-immunolabeling with anti-human ARL13B [Cat. #17711-1-AP]/anti-human rootletin (white/green) in a U-ExM sample of normal canine retina. Scale bars: 1 μm, after correction for expansion factor. **(G-I)** Confocal images of expanded adult canine photoreceptors labeled with anti-human ARL13B [Cat. #17711-1-AP]/anti-AcTub (white/magenta, G, H) and anti-mouse ARL13B [clone N295B/66]/anti-AcTub (white/magenta, I). Rod and cone photoreceptors were identified by labeling with rhodopsin (yellow, G, I) and CAR (cyan, H), respectively. Scale bars: low-magnification view = 1 μm; high-magnification view = 500 nm, after correction for expansion factor. Abbreviations: CC, connecting cilium; IS, inner segment; ONL, outer nuclear layer; PSC, photoreceptor sensory cilium.

We found that ARL13B, a small GTPase belonging to the Arf-like Ras superfamily, is another protein located in the ciliary rootlet of canine photoreceptors using the anti-human ARL13B antibody (17711-1-AP). In studies of the primary cilium, ARL13B has frequently been employed as a marker for the ciliary membrane [48, 49]. Since mutations in *ARL13B* gene have been associated with Joubert syndrome, a systemic ciliopathy accompanied by retinal dystrophy, the localization of ARL13B in photoreceptors has been discussed in various studies, with variable conclusions [50–53]. Although several recent reports using mouse retina have claimed that endogenous ARL13B localizes to the photoreceptor OS (**Supplementary Table 5**), we observed specific localization of ARL13B to the ciliary rootlet in U-ExM-processed canine photoreceptors using the 17711-1-AP antibody (**Figure 8F**).

To ensure that this unique ARL13B localization was not an artifact of the U-ExM process, we performed similar immunostaining on non-expanded cryosections. In non-expanded canine retina, ARL13B localized primarily to the photoreceptor IS, consistent with the results in the U-ExM samples (**Supplementary Figure 7A**). Moreover, we confirmed the specificity of this antibody by using the antigen peptide, which completely blocked ARL13B labeling in the IS (**Supplementary Figure 7C**). Thus, the rootlet-specific labeling with the 17711-1-AP antibody is unlikely to result from nonspecific binding. Similar to rootletin, ARL13B labeled with 17711-1-AP exhibited straight localization in rods and curvilinear localization in cones (**Figure 8G and H**). However, when we tested another anti-ARL13B antibody (clone N295B/66), commonly used in mouse retina studies, we observed OS-specific localization of ARL13B, consistent with previous reports, in both U-ExM and non-expanded samples (**Figure 8I, Supplementary Figure 7B**). The fact that these two widely used anti-ARL13B antibodies exhibited completely different labeling patterns in the canine retina underscores the need for further detailed studies on ARL13B distribution in photoreceptors.

### Molecular architecture of the calyceal processes in canine photoreceptors

Finally, we investigated the localization of protocadherin-15 (PCDH15), an Usher 1 protein, to determine the presence or absence of calyceal processes in canine photoreceptors. Calyceal processes are microvilli-like structures characterized by the localization of Usher proteins in the apical region of the IS of human and nonhuman primate (NHP) photoreceptors and are absent in mouse photoreceptors [24]. Recent research on NHP photoreceptors using expansion microscopy has described the structural organization of the calyceal processes based on the distribution of PCDH15 [17]. We followed this approach on our canine retinal tissues. First, we determined that PCDH15 is located between the OS and IS in both rods and cones in non-expanded retinal cryosections (**Figure 9A**). In the canine rod photoreceptors processed by U-ExM, PCDH15 staining exhibited a characteristic C-shape, and its distribution was limited to the base of the OS (**Figure 9B, Supplementary Movie 3**). In contrast, PCDH15 formed a microvilli-like structure surrounding the OS near the OS/IS junction in the canine cones (**Figure 9C, Supplementary Movie 4**). The distribution pattern of PCDH15 was identical between S-cones and L/M-cones (**Figure 9D and E**). These differences of PCDH15 distribution between rods and cones were consistent with the previous observations in NHP photoreceptors [17]. Additionally, we mapped Whirlin (USH2D), an Usher 2 protein, which localized to the basal side of PCDH15, along with the CC tubulin axoneme in both canine rods and cones. Whirlin exhibited a shorter distribution in cones compared to rods correlating with the difference of CC length between these subtypes (**Figure 9F**). We also attempted to localize another Usher 2 protein, VLGR1 (USH2C), in canine photoreceptors. However, identification of specific VLGR1 signals using a commercially available primary antibody was unsuccessful.

**Figure 9.**
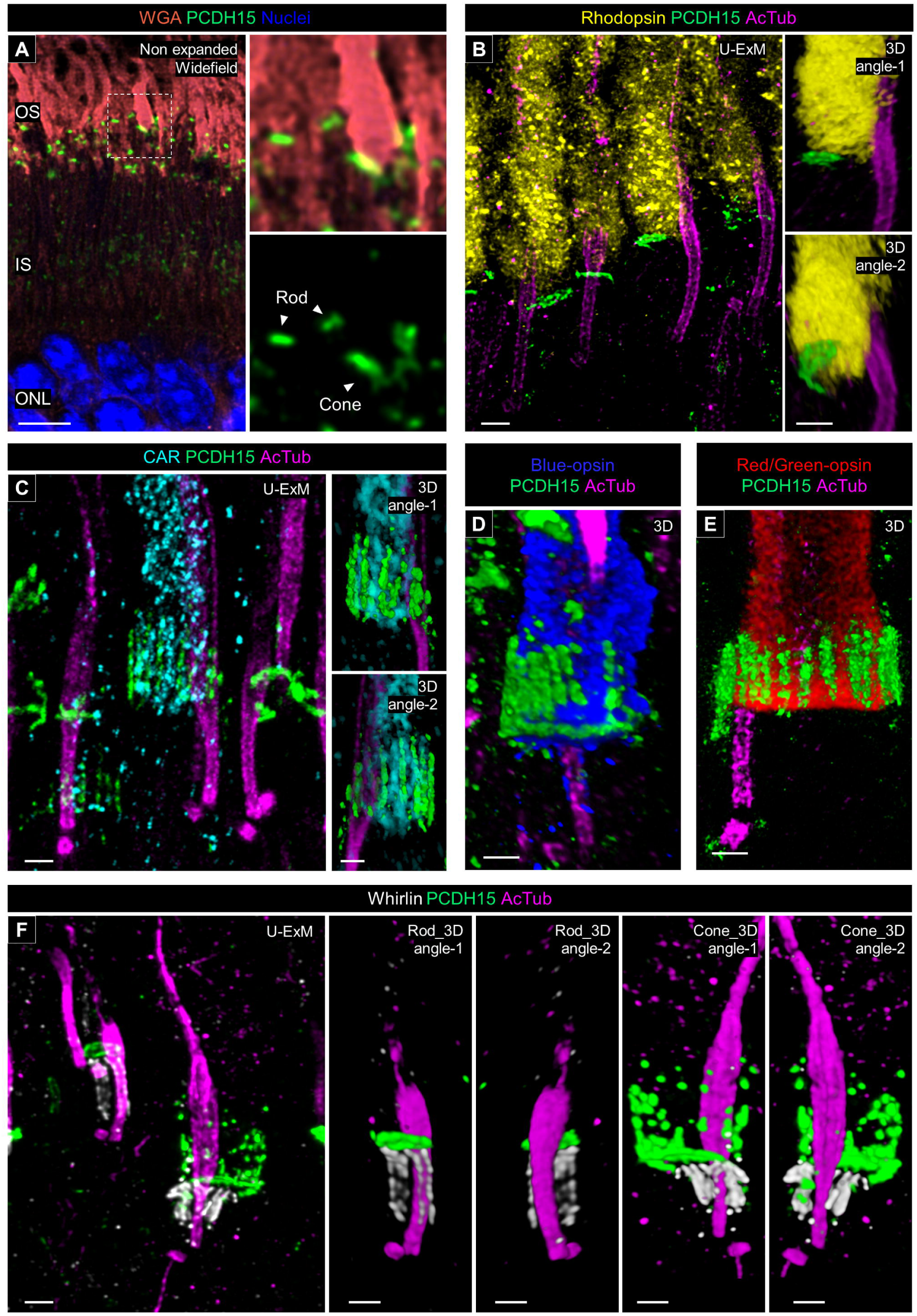
Molecular architecture of calyceal processes in normal canine photoreceptors. **(A)** Widefield view of a non-expanded retinal cryosection stained for wheat germ agglutinin (WGA; orange), and protocadherin 15 (PCDH15; green). The right panels show details of PCDH15 location in rod and cone photoreceptors. Scale bar: 5 μm, without correction for expansion factor. **(B-E)** Confocal images of expanded adult canine photoreceptors stained for PCDH15 (green) and AcTub (magenta). Rod and cone photoreceptors were identified by labeling with rhodopsin (yellow, B), CAR (cyan, C), blue-opsin (blue, D), and red/green-opsin (red, E) respectively. The right panels in (B, C) and (E, D) are showing 3D rendering images of one photoreceptor. Scale bars: 500 nm, after correction for expansion factor. Abbreviations: IS, inner segment; ONL, outer nuclear layer; OS, outer segment.

Finally, we localized actin filaments by immunolabeling of β-actin. In non-expanded canine retina, β-actin was distributed around the IS/OS interface in both cones and rods, although the signal intensity in rods was lower than in cones (**Figure 10A**). However, in U-ExM samples, no actin filaments binding to PCDH15 were observed in canine rods, whereas canine cones displayed distinct actin filaments linking to the PCDH15-labeled microvilli-like architecture (**Figure 10B**). These findings were further supported by Espin labeling, which indicated that calyceal processes-associated actin fibers in canine rods are more fragile than in cones, making them unable to maintain their architecture after expansion (**Figure 10C and D**). Taken together, these results suggest that in canine photoreceptors, while the localization of Usher proteins at the IS/OS junction is present in both rods and cones, the microvilli-like distribution of Usher 1 protein is found only in cone photoreceptors, similar to what is observed in NHP photoreceptors.

**Figure 10.**
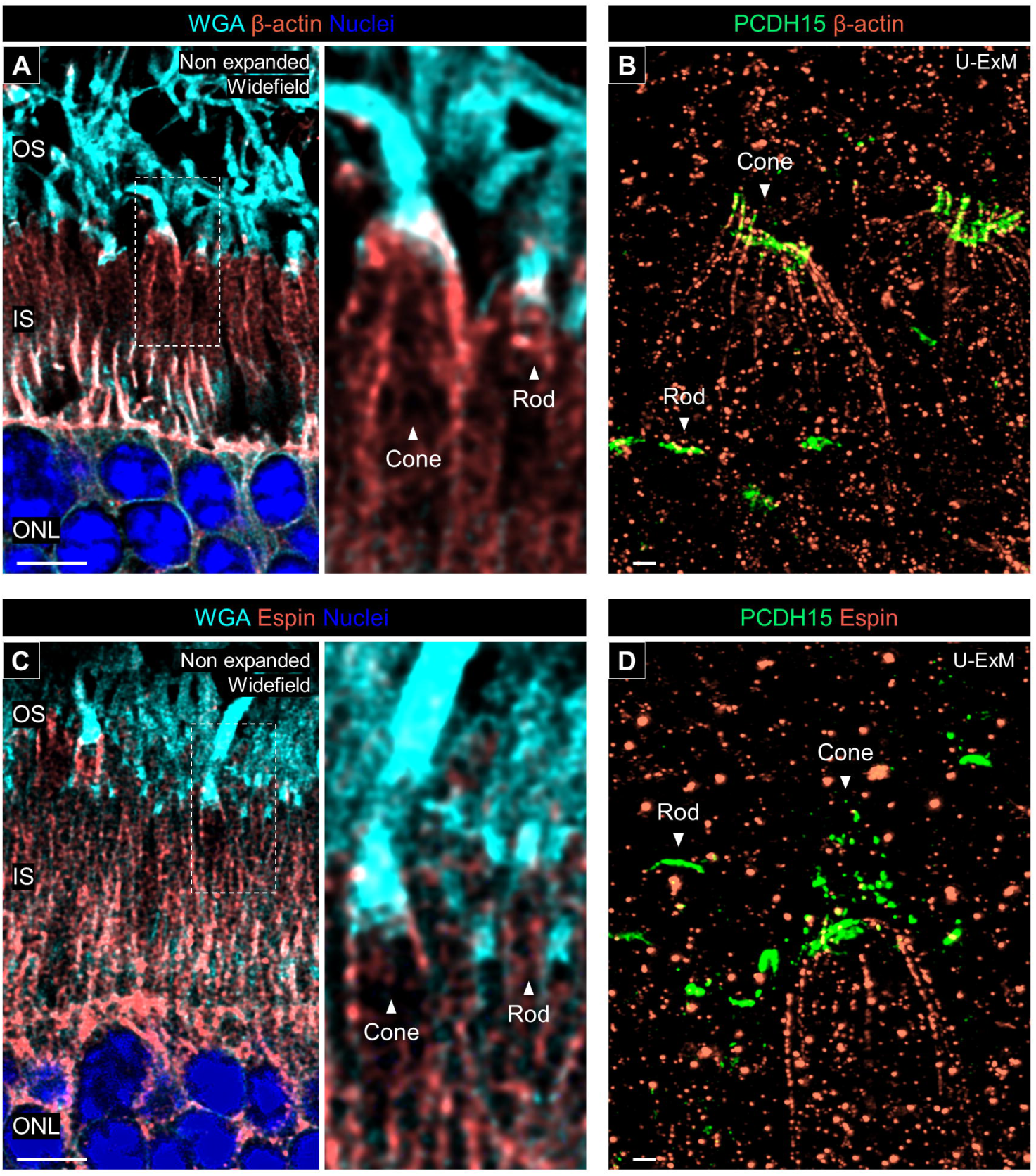
Actin filaments associated with calyceal processes. **(A,C)** Co-labeling with β-actin/WGA (orange/cyan, A) and Espin/WGA (orange/cyan, C) in non-expanded canine retinal tissues. The right panels show details of actin filaments location associated with calyceal processes in rod and cone photoreceptors. Scale bar: 5 μm, without correction for expansion factor. **(B, E)** Co-labeling with β-actin/PCDH15 (orange/green, B) and Espin/PCDH15 (orange/green, D) in U-ExM samples of normal canine retina. Abbreviations: IS, inner segment; ONL, outer nuclear layer; OS, outer segment. Scale bars: 500 nm, after correction for expansion factor.

## Discussion

In the present study, we tested a U-ExM protocol to characterize the molecular architecture of the PSC using non-frozen and frozen archival canine retinal tissues. Utilizing archival samples, particularly from large animals such as dogs and NHP, presents significant advantages in terms of cost efficiency and biological resource management. This approach is not only economical but also allows for the maximization of previously collected biological materials. However, as highlighted in the publication reporting the original U-ExM protocol, the type and duration of fixation can influence the degree of tissue expansion [19]. Earlier studies applying U-ExM to the mouse retina typically used a short duration (15 minutes) of 4%PFA fixation [18, 21]. Here, we demonstrated that the duration of PFA fixation and long-term storage at −80°C do not significantly impact the degree of retinal tissue expansion, achieving approximately 4x isotropic expansion (**Figure 2**). This observation aligns with the recent reports that applied U-ExM to PFA-fixed frozen and formalin-fixed paraffin-embedded mouse tissues [20, 28]. Notably, our results showed that U-ExM enabled super-resolution imaging of archival samples that had been cryopreserved for over a decade, providing comparable imaging quality to that obtained from non-frozen samples. While some physical damage to the OS membrane architecture was observed in the frozen section-derived U-ExM samples compared to the non-frozen samples, the tubulin axoneme structure remained unaffected across the sample processing conditions. Additionally, longer durations of PFA fixation (≥ 24 hours), which are routinely used for cryopreservation of ocular tissues, proved beneficial in preserving the structure of the photoreceptor OS in U-ExM. These findings underscore the robustness of U-ExM for use with archival samples, facilitating ultrastructural observation in large animal tissues where non-frozen samples are often challenging to procure.

In the molecular mapping of the PSC in normal adult canine retinal tissue, we discovered that the tubulin-based PSC architecture differs significantly between the rods and cones (**Figure 3**). Specifically, cones exhibited longer BB and DC, shorter CC, and a longer and wider bulge region compared to rods. Additionally, the tubulin axonemes labeled by anti-AcTub were more prominent in the distal region of cones, which may suggest a more robust axoneme structure in cones than in rods. These structural differences were corroborated by the distinct localization patterns of proteins within each PSC compartment (**Figure 4-7**). While detailed comparisons of PSC molecular architecture between rods and cones have been limited, our findings align with the known variations in the structural features of the OS between these photoreceptor subtypes [54]. Previous studies have consistently reported that the CC length in mature mouse rods ranges from approximately 1100 to 1500 nm [5, 18, 21], which is consistent with the CC-specific protein signal lengths of approximately 1200 to 1400 nm observed in mature canine rods in this study. Similarly, the DC length in mature mouse rods is approximately 300 nm [5], closely matching the 270 nm measured in mature canine rods, indicating a conserved architecture of the CC and BB in rods across these species. However, comparisons of cone PSC architecture between canine and other vertebrates remain challenging due to a paucity of studies investigating the ultrastructure of cone PSC to date. The significant architectural differences between rod and cone PSC observed in this study underscore the necessity for further comprehensive investigations using retinal tissues from a variety of vertebrate species. These investigations are crucial to determine whether the architectural distinctions in the PSC between photoreceptor subtypes are unique to canine or represent a broader biological phenomenon.

An interesting question raised by the observed architectural variation between rod and cone PSC is its involvement in the differential pathological progression among photoreceptor subtypes. A notable group of retinal ciliopathies, known as cone-rod dystrophy (CRD), is characterized by the progressive degeneration of cones preceding that of the rods. In canine models of naturally-occurring retinal ciliopathy, homozygous variants in genes such as *NPHP4*, *NPHP5*, and *RPGRIP1* have been identified as causes of CRD [55–57], despite these genes being similarly expressed in both rods and cones. Previous studies have suggested that these CRD-associated proteins are localized to the CC in photoreceptors [11, 58]. Our current study confirms that NPHP5 and RPGRIP1 are indeed localized outside the tubulin axoneme at the CC in both photoreceptor subtypes (**Figure 6**). Given their comparable distribution, these proteins are likely to have similar roles related to Y-links in both photoreceptor subtypes. Notably, the CC, where the tubulin axoneme of the PSC is anchored to the plasma membrane by Y-links, is significantly shorter in cones than in rods. This structural difference in the CC suggests that cones may be more susceptible to mutations in Y-link-associated genes compared to rods. The relationship between the heterogeneity of ciliopathy-associated phenotypes and the structural variations between rods and cones will be further investigated in future studies, which will involve integrating respective retinal ciliopathy models with U-ExM techniques to elucidate these differences.

Our study has also revealed significant differences in the architecture of the PSC rootlet between rods and cones (**Figure 8**). Using U-ExM, we observed a linear shape of the rootlet in canine rods, which is consistent with the morphology of the rootlet in mouse rods, as previously described in transmission electron photomicrographs [10]. A new finding in this study is the identification of a rootlet branch extending from the basal part of the PSC to the middle of the CC in rods. This branching structure has also been observed in recent transmission electron photomicrographs of canine photoreceptors, where it appears to be positioned on the opposite side of the PSC axoneme across the ciliary pocket [59], although its functional role remains unclear. Another interesting discovery is the unique elliptical shape of the cone rootlet. A recent study in NHP cones suggests that the cone rootlet has a markedly different shape from the commonly recognized form in rod rootlet, although detailed morphological descriptions have been lacking [17]. Here, we have demonstrated a curvilinear rootlet architecture in canine cones using U-ExM. Additionally, unlike the rod rootlet, which extends towards the external limiting membrane, the cone rootlet is entirely contained within the photoreceptor IS. Both rootlet structures likely play a critical role in providing stable intracellular support for the PSC and their associated OS structures. However, further studies are needed to elucidate the biological significance of these morphological differences between rod and cone photoreceptors.

During the mapping of various cilia-associated proteins using U-ExM, we identified several proteins with localization patterns that differed significantly from those reported in previous studies of the vertebrate retina. Earlier studies, based on immunolabeling of non-expanded retinal samples, suggested that CCDC66 localizes to the IS or OS [29–31]. However, given the properties of CCDC66 as a microtubule-binding protein and previous findings from cell line studies [32], its localization to the OS or IS in photoreceptors seems unlikely. Using U-ExM, we found that two anti-CCDC66 antibodies (PA5-60642 and sc-102418) produced highly specific signals for the PSC axoneme, despite showing less specificity in non-expanded retinal tissue (**Figure 4 and Supplementary Figure 4**). The PA5-46125 antibody, which did not detect signals in U-ExM, has a lower homology between its epitope and the corresponding canine CCDC66 protein region compared to PA5-60642 (74% vs. 85%, respectively), suggesting that this might affect its ability to accurately label the target protein. The improved specificity of the other anti-CCDC66 antibodies (PA5-60642 and sc-102418) for the PSC axoneme could be attributed to the enhanced accessibility of epitopes following the U-ExM procedure. On the other hand, further investigation is required to explain the rootletin-specific signals obtained with the anti-human ARL13B antibody (17711-1-AP). The labeling patterns of the two anti-ARL13B antibodies used in this study were consistent between non-expanded and U-ExM samples, although the patterns differed between the antibodies (**Figure 8 and Supplementary Figure 7**). A recent study using ARL13B-mCherry Centrin2-GFP double transgenic mice reported an IS-specific and rootlet-like ARL13B-mCherry distribution, which closely mirrors our findings in the canine retina, albeit restricted to the first 14 days after birth [60]. Moreover, given that ARL13B-mcherry was widely detected from the OS to IS of mature retina in the same transgenic mice, it is challenging to completely dismiss the rootlet-specific labeling of the 17711-1-AP antibody observed in this study.

Finally, the application of U-ExM provided novel insights into the architecture of calyceal processes in canine photoreceptors (**Figure 9 and 10**). Our findings indicate that canine cones possess microvilli-like structures characterized by the distribution of PCDH15 around the basal part of the OS and the presence of associated actin filaments. In contrast, canine rods lack microvilli-like PCDH15 distribution, despite the localization of PCDH15 at the IS/OS junction. These differences in PCDH15 distribution between rods and cones are similar to those observed in photoreceptors of NHPs in a previous study [17]. Interestingly, at the electron microscopy level, microvilli-like structures have been observed in NHP rods, though in smaller numbers compared to cones [24]; however, recent studies have shown an absence of microvilli-like PCDH15 distribution in these rods [17]. This might suggest that the distribution of PCDH15 in rods does not align with the microvilli-like architecture considered as calyceal processes. To conclusively determine the presence or absence of calyceal processes in canine rods, further detailed morphological studies using electron microscopy are necessary. The localization of Usher proteins (PCDH15 and Whirlin) as well as actin filament-related proteins in canine photoreceptors aligns with the patterns reported in NHP photoreceptors, suggesting that a canine IRD model may provide valuable insights into pathogenesis of Usher syndrome, for which effective pathological models are challenging to develop in rodents.

In conclusion, our study highlights the effectiveness of super-resolution imaging using U-ExM to characterize the molecular architecture of the canine PSC. By employing U-ExM, we have achieved detailed molecular mapping of numerous retinal ciliopathy-related proteins and have identified significant structural differences in the PSC between rods and cones in canine photoreceptors (**Figure 11**). These insights gained from this study pave the way for a better understanding of the alterations in PSC molecular architecture and the efficacy of AAV-based gene augmentation therapies in established canine models of retinal ciliopathy, such as those with mutations in *RPGR* [61], *NPHP5* [56], *RPGRIP1* [57], and *CCDC66* [31], using U-ExM.

**Figure 11.**
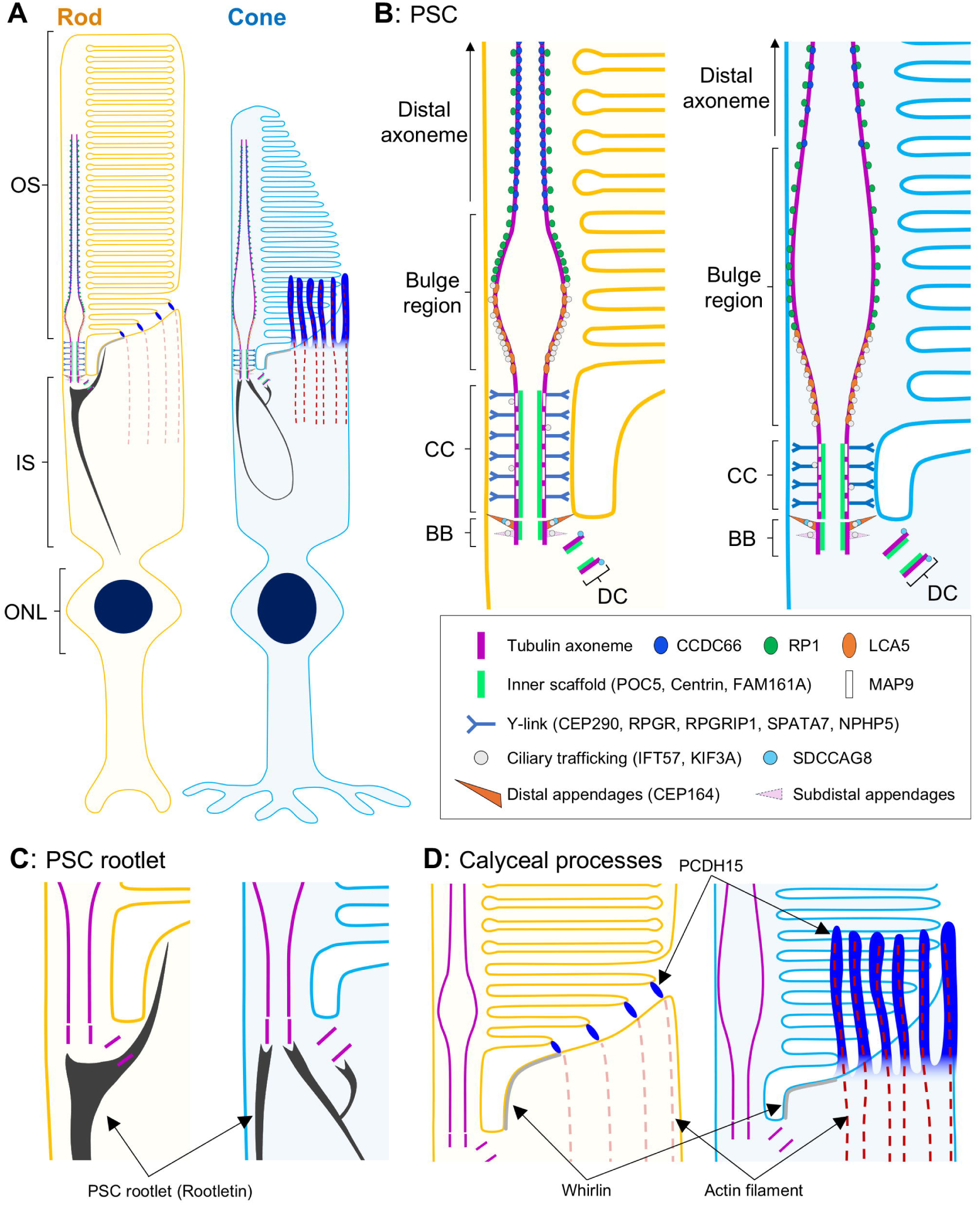
Schematic summary diagrams of the molecular architecture of PSC and calyceal processes, and the distribution of retinal ciliopathy-associated proteins in normal canine rod and cone photoreceptors. **(A)** Overview of the architecture of normal canine rod (left) and cone (right) photoreceptors. **(B)** Comparison of PSC molecular architecture between rod and cone photoreceptors. Cone PSC has a longer and wider bulge region, a longer BB, a longer DC, and a shorter CC than those of rod PSC. **(C)** Structural differences of PSC rootlet between rod and cone photoreceptors. Rod photoreceptors have a straight rootlet extending toward the ONL, whereas the rootlet of cones forms a curvilinear shape. **(D)** Distribution of calyceal processes-related proteins in rod and cone photoreceptors. In rod photoreceptors, the localization of PCDH15 is limited to the IS/OS junction, while in cone photoreceptors, PCDH15 shows a microvilli-like distribution accompanied by actin filaments. Abbreviations: BB, basal body; CC, connecting cilium; DC, daughter centriole; IS, inner segment; ONL, outer nuclear layer; OS, outer segment; PSC, photoreceptor sensory cilium.

### Material and methods

### Animals

Retinal tissues from normal adult dogs were used for this study (**Supplementary Table 1**). All dogs were housed under identical conditions, including diet and ambient illumination with a 12-hour light/12-hour dark cycle, at the Retinal Disease Studies Facility (RDSF) at the University of Pennsylvania. The studies strictly adhered to the ARVO Statement for the Use of Animals in Ophthalmic and Vision Research and were approved by the Institutional Animal Care and Use Committee of the University of Pennsylvania.

### Tissue processing for long-term cryopreservation

The procedures for long-term cryopreservation of normal adult canine ocular tissues were performed as described in previous publications [62, 63]. In brief, following enucleation of euthanized dogs, the posterior eyecups of the globes were isolated and the vitreous was gently extracted ∼ 1 hour after fixation in 4% PFA in 0.1 M phosphate-buffered saline (PBS). Fixation of the posterior eye cups was pursued for an additional 2 hours, followed by an additional 24-hour fixation in 2% PFA in 0.1 M PBS at 4°C. Subsequently, the eyecups were dissected into four quadrants, and the tissues were sequentially cryoprotected for 24 hours each in 15% and 30% sucrose solutions in 0.1 M PBS at 4°C. Following cryopreservation, the dissected tissues were embedded in optimal cutting temperature (OCT) medium within disposable molds. The molds were then placed in shallow metal cups containing methylbutane and immersed in a bath of liquid nitrogen for rapid freezing. The frozen OCT blocks were stored at −80°C until cryosectioning was conducted. All cryopreserved samples used in this study had been stored for more than 12 years at −80°C (**Supplementary Table 1**).

### Immunohistochemistry on non-expanded cryosectioned retinal tissues

The frozen OCT blocks were cryosectioned at a thickness 10 µm and stored at −20°C until further processing. Detailed methodology for immunostaining of non-expanded canine retinal cryosection has been described in a previous publication [34]. Frozen sections were dried at room temperature (RT) for 60 minutes and then incubated in Dulbecco’s PBS (D-PBS) for 10 minutes to remove the OCT compound. Antigen retrieval was performed by treating the sections with pepsin reagent for 1 minute, followed by incubation with blocking buffer (4.5% cold fish gelatin, 0.1% sodium azide, 5% bovine serum albumin, and 0.25% Triton X-100 in D-PBS) for 60 minutes at RT. Retinal sections were then incubated overnight at 4°C with primary antibodies (**Supplementary Table 3**) diluted in blocking buffer. In the specificity confirmation study of the anti-human ARL13B antibody (Cat. #17711-1-AP), 50 μg/mL of ARL13B fusion protein (Cat. #Ag12015) was pre-incubated with the antibody at RT for 2 hours before being applied to retinal tissue. After three washes with D-PBS containing 0.1% of tween20 (PBS-T), antigen-antibody complexes were visualized with Alexa Fluor-conjugated secondary antibodies (**Supplementary Table 4**) and counterstained using Hoechst 33342. Fluorescein-conjugated Wheat Germ Agglutinin (WGA) was diluted 1:200 in D-PBS and treated for 15 minutes after secondary antibody incubation. The reagents used for conventional immunostaining were listed in Supplementary Table 2.

### Ultrastructure Expansion Microscopy (U-ExM) on canine retina

Based on a U-ExM protocol previously developed for mouse retina [18], we tested U-ExM on non-frozen and frozen archival canine retinal tissues. A schematic overview, photographs of each step, and reagents associated with the U-ExM protocol used in this study are summarized in Supplementary Figure 1 and 2 and in Supplementary Table 2.

#### (a) U-ExM on non-frozen canine neuroretinas

After enucleation, the anterior part of eyeball, including the cornea, lens, and vitreous, was gently removed. The posterior eyecups were fixed in PFA solution for either short (4% PFA for 15 minutes) or long (4% PFA for 3 hours followed by 2% PFA for 24 hours) durations, and then 5 mm punches were taken from the eyecups using a disposable biopsy punch (**Supplementary Figure 2A**). Once isolated from RPE-choroid-sclera complex (**Supplementary Figure 2B, 2C**), a 5 mm retinal punch was placed in a 14-mm microwell of a 35-mm petri dish and then incubated overnight in 300 μL of 2% acrylamide (AA) + 1.4% formaldehyde (FA) solution at 37°C.

After removing the AA/FA solution, 45 μL of monomer solution (MS) composed of 25 μL of sodium acrylate (SA; stock solution at 38% [w/w] diluted with nuclease-free water), 12.5 μL of AA (40% solution), 2.5 μL of N,N′-methylenebisacrylamide (BIS; 2% solution), and 5 μL of 10× PBS was added to the microwell. After 90 minutes of incubation at RT, the MS was removed and 90 μL of MS containing ammonium persulfate (APS) and tetramethylethylenediamine (TEMED) at a final concentration of 0.5% was dropped onto the tissue, followed by covering with a 12 mm circular cover glass (**Supplementary Figure 2D**). Subsequently, the 35-mm petri dish containing the tissue and MS solution was placed on ice for 45 minutes, followed by 60 minutes incubation at 37°C to allow gelation.

Next, 1 mL of denaturation buffer (200 mM Sodium Dodecyl Sulfate (SDS), 200 mM Sodium Chloride (NaCl), 50 mM Tris Base in deionized distilled water (ddH_2_O), pH 9) were added to the petri dish for 15 minutes at RT with gentle agitation. The gel was carefully detached from the microwell using a spatula and then incubated in a 1.5 mL tube filled with denaturation buffer for 90 minutes at 95°C. Immediately after the denaturation step, the gel was transferred to a beaker filled with ddH_2_O, and the ddH_2_O was replaced twice, every 30 minutes, for the first-round of expansion.

The following day, after measuring the diameter with a ruler (**Supplementary Figure 2E**), the expanded gel containing retinal tissue was sliced with a razor blade to obtain approximately 1-2 mm thick cross sections. These sections were then placed in a well of 24-well plate filled with D-PBS containing calcium & magnesium (D-PBS-Ca^2+^Mg^2+^) for the immunostaining steps.

#### (b) U-ExM for cryosections from frozen retinal tissue

Twenty-μm-thick cryosections were obtained from the frozen posterior cup tissues that had been fixed with PFA using the longer protocol, cryoprotected in sucrose, and embedded in OCT media. The frozen sections on a slide glass were thawed at RT for 10 minutes (**Supplementary Figure 2F**), and a square was drawn around the tissue section using a hydrophobic marker (ImmEdge Pen). After drying for 60 minutes at RT, 100 µL of D-PBS-Ca^2+^Mg^2+^ was added into the hydrophobic square for 10 minutes to wash out the OCT compound.

After removal of D-PBS, the sections were reacted with the AA/FA solution overnight, followed by penetration with MS for 90 minutes, as was done for non-frozen retinal punches. For each retinal section, 45 µL of MS containing APS and TEMED per section was dropped onto the tissue, and a 12 mm circular cover glass was placed on the solution (**Supplementary Figure 2G**). After performing gelation, denaturation, and the first-round expansion as done for non-frozen retinal punches, the diameter of the gel was measured with a ruler (**Supplementary Figure 2H**). The gel containing the retinal tissue was then cut into approximately 1 cm squares using a razor blade and then in a well of 24-well plate filled with D-PBS-Ca^2+^Mg^2+^.

#### Immunohistochemistry for U-ExM samples

After 10 minutes of incubation of the gels in D-PBS, the solution in the well was replaced with 300 μL of blocking buffer containing the primary antibodies (**Supplementary Table 3**). The gels were incubated with the primary antibody solution overnight at RT with gentle shaking, followed by 3 washes with 1 mL of PBS-T for 5 minutes each. Alexa Fluor-conjugated secondary antibodies (**Supplementary Table 4**) were diluted in the blocking buffer at a concentration of 1/500 and incubated with the gel for 2 hours at RT, protected from light. Next, 100 µL of Hoechst 33342 solution, diluted 1/1000 in D-PBS, was added to the wells and gently shaken for 30 minutes, followed by 3 washes with 1 mL of PBS-T. After the washes, each gel was transferred using a spatula to a well of a 6-well plate filled with 10 mL ddH_2_O and incubated for 30 minutes to achieve the second-round expansion. The ddH_2_O was replaced twice, every 30 minutes. The gels were then stored in ddH_2_O at 4°C, protected from light, until confocal imaging.

### Confocal imaging

Prior to imaging, 14-mm microwells of 35-mm petri dishes were filled with 0.1 mg/mL poly-D-lysine and kept at RT for 1 hour for coating to prevent drifting of the gels. After removing the poly-D-lysine solution, the microwells were washed three times with ddH_2_O, dried, and then stored at 4°C.

Image acquisition was performed on the Leica Stellaris 8 FALCON confocal FLIM microscope using a 63× (1.20 NA) water objective. The lateral and axial views were obtained by orienting the expanded gel on the bottom of the microwell to face either the retinal cross-section or photoreceptor outer segment downward. Z-stacks for the lateral view were acquired with 0.2 μm z-intervals, an x, y pixel size of 45.09 nm, and an imaging speed of 400 Hz. Representative images of the lateral view are shown as maximum projection images unless specified. Sigle images of the axial view were obtained with an x, y pixel size of 22.55 nm and an imaging speed of 100 Hz. The images were processed using Leica Application Suite X (LAS X) software for deconvolution with LIGHTNING function, as well as for building merged and maximum projection images, and 3D rendering.

### Quantification

#### Gel diameter

The gel diameter after the first-round expansion was measured using a ruler and then divided by the gel diameter before expansion (1.2 mm) to calculate the degree of gel expansion.

#### Expansion factor

The expansion factor was calculated by dividing the width of the AcTub-labeled axoneme on the CC of U-ExM samples by the previously reported CC diameter of non-expanded mice photoreceptors (200.84 nm), quantified using cryo-electron tomography [5]. A total of 40 photoreceptors in all U-ExM conditions were subjected to measurement of AcTub signal width using LAS X software manually. The width of the CC was consistent across all U-ExM conditions, and the expansion factor in this study was determined to be 3.83 by averaging the values of all U-ExM conditions. This expansion factor was used for every quantification.

#### Protein signal length, width, and distance from the proximal end of the BB

Lengths, widths, and distance of protein signals were measured using the “Draw scalebar” function in LAS X software manually to fit the photoreceptor curvature. AcTub-labeled axoneme spread was measured at four different locations of the rod and cone photoreceptors corresponding to the CC or bulge region: +1000, +500, 0, and −500 nm. The distal end of the glutamylation signal located outside the tubulin axoneme was used to define the 0 location. Each measurement was subsequently corrected using the expansion factor.

### Statistical analysis

Comparisons between 2 groups were performed using the nonparametric Mann-Whitney U test. Comparisons among more than 3 groups were conducted using the nonparametric Kruskal-Wallis test with Dunn’s multiple-comparison test. Every measurement was performed on at least 2 individual eyes. Quantitative data are represented as scatter dot plots, with the center line and error bars in the graphs indicating the mean and standard deviation (± SD), respectively. The significance level is denoted as follows: nonsignificant (n.s.) P > 0.05, *P < 0.05, **P < 0.01, ***P < 0.001, ****P < 0.0001. All statistical analyses were performed using GraphPad Prism 10. The data underlying the graphs shown in all the figures are included in Supplementary Data.

### Data availability

All data are available in the main text or the supplementary information.

## Author Contributions

K.T. and W.A.B. participated in the conception, experimental design, and interpretation of the data for this work. K.T. was responsible for the acquisition and analysis of data, statistical analysis, and the drafting of the manuscript. R.S. and W.A.B. reviewed and edited the manuscript. All authors reviewed the manuscript, approved its submission, and take full responsibility for the manuscript.

## Acknowledgements

We express our gratitude to Qin Liu and Eric A. Pierce of Harvard Medical School for providing the RP1 antibody [64]; Penn Vet Imaging Core Facility (Research Resource Identifier: SCR_022764) for technical support; the staff of the Division of Experimental Retinal Therapies for their excellent support; and the staff of RDSF at University of Pennsylvania for their outstanding animal care. This study was supported by NIH grants R01EY006855, R01EY017549, R01EY033049, P30EY001583, S10OD032305-01A1, and Foundation Fighting Blindness.

