## Supplemental tables for "Mapping protein distribution in the canine photoreceptor sensory cilium and calyceal processes by ultrastructure expansion microscopy"

**Supplementary Table 1. Normal adult canines used in this study.**

| Dog ID | Eye | Sex | DOB | DOD | Age (months) | Storage at -80°C |
|---|---|---|---|---|---|---|
| **Fresh retina fixed using short duration** | | | | | | |
| 1180608 | OS | M | 4-May-2023 | 14-Sep-2023 | 4 | n/a |
| RC724 | OD | M | 9-Jul-2022 | 28-Sep-2023 | 14 | n/a |
| RC724 | OS | M | 9-Jul-2022 | 28-Sep-2023 | 14 | n/a |
| **Fresh retina fixed using long duration** | | | | | | |
| 1180560 | OD | M | 4-May-2023 | 12-Sep-2023 | 4 | n/a |
| 1180560 | OS | M | 4-May-2023 | 12-Sep-2023 | 4 | n/a |
| **Frozen archival retina** | | | | | | |
| GI97 | OD | F | 15-Apr-2010 | 29-Nov-2010 | 7 | > 12 years |
| GI97 | OS | F | 15-Apr-2010 | 29-Nov-2010 | 7 | > 12 years |
| RC667 | OD | F | 17-Mar-2010 | 3-Nov-2010 | 7 | > 12 years |
| RC667 | OS | F | 17-Mar-2010 | 3-Nov-2010 | 7 | > 12 years |

DOB, day of birth; DOD, day of death; F, female; M, male; OD, right eye; OS, left eye.

**Supplementary Table 2. Reagents and materials used in this study.**

| Product | Supplier | Reference |
|---|---|---|
| **Tissue processing and storage** | | |
| 20% paraformaldehyde solution | Electron microscopy sciences | 15713-S |
| Sterile Disposable Biopsy Punch 5mm | Sklar | 96-1120 |
| Sucrose | Fiser chemical | S2-500 |
| Tissue-Tek O.C.T. Compound | Sakura finetek | 4583 |
| Tissue Embedding Disposable Molds | EBSciences | H1513 |
| **U-ExM** | | |
| 14-mm microwell/35-mm petri dish | MatTek | P35G-1.5-14-C |
| 12 mm Circular Cover Glasses | Fisher Scientific | 12541001 |
| Acrylamide | Sigma-Aldrich | A4058 |
| Ammonium Persulfate (APS) | Bio-Rad | 1610700 |
| Formaldehyde solution | Sigma-Aldrich | F8775 |
| Nuclease-Free Water | Invitrogen | AM9937 |
| ImmEdge Pen | Vector laboratories | H-4000 |
| N, N′-methylenbisacrylamide (BIS) | Sigma-Aldrich | M1533 |
| Phosphate Buffered Saline (PBS), 10x | Bio-Rad | 161-0780 |
| Poly-D-Lysine | Gibco | A3890401 |
| Sodium Acrylate (SA) | Sigma-Aldrich | 408220 |
| Sodium Chloride (NaCl) | Fisher Chemical | S271-3 |
| Sodium Dodecyl Sulfate (SDS) | Fisher Chemical | BP166-500 |
| Tetramethylethylenediamine (TEMED) | Bio-Rad | 161-0800 |
| Tris Base | Fisher Chemical | BP152-5 |
| **Immunohistochemistry** | | |
| Bovine Serum Albumin | Sigma-Aldrich | A7906 |
| D-PBS with calcium & magnesium | Corning | 21-030-CM |
| Gelatin from cold water fish skin | Sigma-Aldrich | G7765 |
| Hoechst 33342 | Thermo Scientific | 62249 |
| Pepsin Reagent, Antigen Retriever | Sigma-Aldrich | R2283 |
| Sodium Azide | Sigma-Aldrich | S8032 |
| Triton X-100 | Fisher Scientific | BP151 |
| Tween20 | Bio-Rad | 170-6531 |

**Supplementary Table 3. Primary antibodies used in this study.**

| Target | Host species | Supplier | Reference | Dilution |
|---|---|---|---|---|
| Acetylated α-tubulin | Mouse monoclonal IgG2b | Sigma-Aldrich | T7451 | 1/1000 |
| Acetylated α-tubulin | Rabbit monoclonal IgG | abcam | ab179484 | 1/1000 |
| α-tubulin (non-acetylated) | Rabbit monoclonal IgG | abcam | ab18251 | 1/1000 |
| ARL13B | Rabbit polyclonal IgG | Proteintech | 17711-1-AP | 1/200 |
| ARL13B (clone N295B/66) | Mouse monoclonal IgG2a | abcam | ab136648 | 1/200 |
| β-actin | Rabbit polyclonal IgG | abcam | ab8227 | 1/500 |
| Blue-opsin | Rabbit polyclonal IgG | Sigma-Aldrich | AB5407 | 1/500 |
| CCDC66 | Rabbit polyclonal IgG | Invitrogen | PA5-60642 | 1/200 |
| *CCDC66 | Rabbit polyclonal IgG | Invitrogen | PA5-46125 | 1/200 |
| CCDC66 | Rabbit polyclonal IgG | Santa Cruz Biotechnology | sc-102418 | 1/200 |
| Centrin | Mouse monoclonal IgG2aκ | Sigma-Aldrich | 04-1624 | 1/200 |
| CEP164 | Rabbit polyclonal IgG | Proteintech | 22227-1-AP | 1/200 |
| CEP290 | Rabbit polyclonal IgG | Proteintech | 22490-1-AP | 1/200 |
| Cone arrestin | Goat polyclonal IgG | Custom made (Beltran Lab) | ref. 63 | 1/400 |
| Espin | Mouse monoclonal IgG1κ | Santa Cruz Biotechnology | sc-515657 | 1/200 |
| FAM161A | Rabbit polyclonal IgG | Invitrogen | PA5-56935 | 1/100 |
| γ-tubulin | Mouse monoclonal IgG1 | Sigma-Aldrich | T6557 | 1/1000 |
| Glutamylation (GT335) | Mouse monoclonal IgG1κ | AdipoGen | AG-20B-0020 | 1/1000 |
| IFT57 | Rabbit polyclonal IgG | Novus | NBP1-32932 | 1/200 |
| KIF3A | Rabbit polyclonal IgG | abcam | ab11259 | 1/100 |
| LCA5 | Rabbit polyclonal IgG | Proteintech | 19333-1-AP | 1/200 |
| MAP9 | Rabbit polyclonal IgG | Proteintech | 26078-1-AP | 1/200 |
| NPHP5 | Rabbit polyclonal IgG | Proteintech | 15747-1-AP | 1/200 |
| PCDH15 | Sheep polyclonal IgG | R&D systems | AF6729 | 1/300 |
| POC5 | Rabbit polyclonal IgG | Bethyl | A303-341A | 1/200 |
| Red/Green-opsin | Rabbit polyclonal IgG | Sigma-Aldrich | AB5405 | 1/500 |
| Rhodopsin | Mouse monoclonal IgG1 | Sigma-Aldrich | MAB5316 | 1/1000 |
| Rhodopsin | Rabbit polyclonal IgG | Sigma-Aldrich | AB9279 | 1/1000 |
| Rootletin | Human monoclonal IgG | AbD Serotec | HCA009 | 1/200 |
| RP1 | Chicken polyclonal IgY | Custom made (Liu Lab) | ref. 64 | 1/200 |
| RPGR | Rabbit polyclonal IgG | Proteintech | 16891-1-AP | 1/200 |
| RPGRIP1 | Rabbit polyclonal IgG | Sigma-Aldrich | HPA042955 | 1/200 |
| SDCCAG8 | Rabbit polyclonal IgG | Novus | NBP2-13288 | 1/200 |
| SPATA7 | Rabbit polyclonal IgG | Proteintech | 12020-1-AP | 1/200 |
| * VLGR1 | Rabbit polyclonal IgG | Atlas Antibodies | HPA067503 | 1/100 |
| Whirlin | Rabbit polyclonal IgG | Proteintech | 25881-1-AP | 1/200 |

* No unique signals could be detected in the U-ExM samples.

**Supplementary Table 4. Secondary antibodies and lectin fluorescent conjugates used in this study.**

| Target | Host species | Alexa Fluor | Supplier | Reference |
|---|---|---|---|---|
| Chicken IgY (H+L) | Goat polyclonal IgG | 488 | Invitrogen | A-11039 |
| Mouse IgG (H+L) | Donkey polyclonal IgG | Plus 488 | Invitrogen | A-32766 |
| Mouse IgG2b | Goat polyclonal IgG | 488 | Invitrogen | A-21141 |
| Sheep IgG (H+L) | Donkey polyclonal IgG | 488 | Invitrogen | A-11015 |
| Human IgG (H+L) | Goat polyclonal IgG | 568 | Invitrogen | A-21090 |
| Mouse IgG1 | Goat polyclonal IgG | 568 | Invitrogen | A-21124 |
| Mouse IgG2a | Goat polyclonal IgG | 568 | Invitrogen | A-21134 |
| Mouse IgG (H+L) | Donkey polyclonal IgG | 568 | Invitrogen | A-10037 |
| Rabbit IgG (H+L) | Donkey polyclonal IgG | 568 | Invitrogen | A-10042 |
| Goat IgG (H+L) | Donkey polyclonal IgG | Plus 647 | Invitrogen | A-32849 |
| Mouse IgG1 | Goat polyclonal IgG | 647 | Invitrogen | A-21240 |
| Rabbit IgG (H+L) | Goat polyclonal IgG | 647 | Invitrogen | A-21246 |
| Wheat Germ Agglutinin (WGA) | - | Fluorescein | Vector Laboratories | FL-1201 |

All secondary antibodies were used at a 1/1000 dilution for non-expanded IHC and 1/500 for U-ExM staining protocols.

**Supplementary Table 5. Summary of immunohistological analysis for CCDC66 and ARL13B from previous publications and the current study.**

### CCDC66

| **Retinal tissues** | Mouse, Dog, Human | Mouse | Dog | Dog | | |
|---|---|---|---|---|---|---|
| **Antibodies** | Cat. #sc-102418 | | Cat. #PA5-46125 | Cat. #PA5-60642 | Cat. #PA5-46125 | Cat. #sc-102418 |
| **Epitopes** | Human CCDC66 (*an internal region) | | Human CCDC66 (AA 45-94) | Human CCDC66 (AA 355-449) | Human CCDC66 (AA 45-94) | Human CCDC66 (*an internal region) |
| **Methods** | IHC | IHC | IHC | 1) IHC (U-ExM)<br>2) IHC (no-expansion) | 1) IHC (U-ExM)<br>2) IHC (no-expansion) | 1) IHC (U-ExM)<br>2) IHC (no-expansion) |
| **Distribution in photoreceptors** | IS (**OS) | OS | OS | 1) **Ciliary axoneme**<br>2) IS + CC | 1) No detection<br>2) OS | 1) **Ciliary axoneme**<br>2) No detection |
| **References** | Dekomien G et al. 2010 (ref. 29) | Gerding WM et al. 2011 (ref. 30) | Murgiano L et al. 2020 (ref. 31) | Current study | | |

* Immunogen sequence not provided.
** Faint immunoreactivity in the OS.

### ARL13B

| **Retinal tissues** | Mouse | Mouse | Mouse | Dog | |
|---|---|---|---|---|---|
| **Antibodies** | Custom made | 1) Cat. #17711-1-AP<br>2) Clone N295B/66, Cat. #75-287 | Clone N295B/66, Cat. #75-287 | Cat. #17711-1-AP | Clone N295B/66, Cat. #ab136648 |
| **Epitopes** | Mouse ARL13B (AA 208–428) | 1) Human ARL13B (AA 1-321)<br>2) Mouse ARL13B (AA 208-427) | Mouse ARL13B (AA 208-427) | Human ARL13B (AA 1-321) | Mouse ARL13B (AA 208-427) |
| **Methods** | IHC | IHC | IHC | 1) IHC (U-ExM)<br>2) IHC (no-expansion) | 1) IHC (U-ExM)<br>2) IHC (no-expansion) |
| **Distribution in photoreceptors** | IS | OS (***IS) | OS | 1) IS (Ciliary rootret)<br>2) IS (Ciliary rootret) | 1) OS<br>2) OS |
| **References** | Kim YK et al. 2013 (ref. 51) | Hanke-Gogokhia C et al. 2017 (ref. 53) | Dilan TL et al. 2019 (ref. 52) | Current study | |

*** Partial and limited distribution in the IS.
